# An interferon-independent innate immune response to double stranded RNA in embryonic stem cells

**DOI:** 10.64898/2026.03.24.713854

**Authors:** Pengcheng Ma, Jianlin Xu, Tong Lu, Ronghua Luo, Yuwei Li, Jiangwei Lin, Xinglou Yang, Yongtang Zheng, Ming Shao, Bingyu Mao

**Author notes:** Corresponding author. (M.S.); (B.M.). These authors contributed equally to this work (Pengcheng Ma, Jianlin Xu, Tong Lu).

## Abstract

In vertebrate early embryos and embryonic stem cells, the interferon (IFN)–centered double-stranded RNA (dsRNA) sensing and signaling pathway is markedly suppressed, implying the existence of an alternative repertoire of dsRNA sensors in these stages. We recently reported that dsRNA treatment triggers translation inhibition using Prkra as a sensor in zebrafish and mouse early embryos. Independently, here we show that dsRNA stimulates the expression of a subset of interferon-stimulated genes (ISGs) in the absence of IFN production, establishing a defensive state in mouse embryonic stem cells (mESCs). Upon dsRNA stimulation, the multifunctional DExD/H-box RNA helicase Dhx9 is recruited into dsRNA-induced condensates, where it promotes the recruitment and functional suppression of the Mdm2/Cul4A ubiquitin ligase machinery by excluding their substrate adaptor Ddb1, thereby stabilizing p53 and Stat1. Dhx9, p53 and Stat1 then cooperate to stimulate an ISG response. This signaling is important for the defense against ZIKV infection in mESCs as well as other dsRNA stresses. Such a cascade is conserved in human ESCs, while in zebrafish embryos, p53 but not Stat1 is required for the transcriptional response. Our findings define an immediate, cell-autonomous innate immune pathway operating in ESCs and vertebrate embryos.

---

In differentiated vertebrate cells, double-stranded RNAs (dsRNAs)—whether derived from viral infection or produced endogenously—are detected by pattern recognition receptors (PRRs), initiating an inflammatory response (*1*, *2*). The major cytoplasmic sensors for dsRNA are retinoic acid-inducible gene I (RIG-I) and melanoma differentiation-associated gene 5 (MDA5), while Toll-like receptor 3 (TLR3) acts as a key endosomal dsRNA sensor. Upon binding dsRNA, RIG-I and MDA5 undergo conformational changes and oligomerization, which then activate the mitochondrial antiviral signaling protein (MAVS), leading to the activation of transcription factors interferon regulatory factor 3 (IRF3) and nuclear factor-κB (NF-κB). These factors then drive the expression of type I interferons (IFNs) and proinflammatory cytokines (*3-6*). Secreted type I IFNs further stimulate the expression of IFN-stimulated genes (ISGs) *via* the Janus kinase-signal transducer and activator of transcription (JAK-STAT) pathway, thereby establishing an antiviral state both within infected cell and in neighboring bystander cells (*7*). A central transcription factor in this process is STAT1: upon IFN binding to the IFN-α/β receptor (IFNAR), STAT1 becomes phosphorylated, dimerizes, translocates into the nucleus, and initiates ISG transcription (*4*). Beyond the RIG-I/MDA5 pathway, dsRNA can also directly activate protein kinase R (PKR), which then phosphorylates the eukaryotic translation initiation factor eIF2α, leading to a global suppression of protein synthesis—a conserved antiviral defense strategy (*8-10*).

Notably, this canonical dsRNA response is attenuated in pluripotent stem cells. In both mouse and human embryonic stem cells (mESCs and hESCs), as well as human induced pluripotent stem cells (hiPSCs), MDA5 expression is markedly low. Although RIG-I is expressed, the absence of the adaptor MITA/STING (mediator of IRF3 activation/stimulator of interferon genes) renders these cells unresponsive to dsRNA stimulation (*11*). In mESCs specifically, MAVS expression is further suppressed by microRNAs (miRNAs), effectively inactivating the RIG-I/MDA5–MAVS pathway (*12*). Additionally, ESCs exhibit reduced sensitivity to type I IFNs—a feature thought to be essential for preserving their self-renewal capacity and pluripotent state (*13*). Instead, RNA interference has been suggested to be a major defense mechanism against dsRNA in ESCs, especially during virus infection (*14-16*). However, whether pluripotent cells use distinct dsRNA-sensing mechanisms to drive transcriptional *versus* posttranscriptional responses is only beginning to be elucidated.

Our recent work revealed that dsRNAs trigger Prkra (protein kinase, interferon-inducible double-stranded RNA dependent activator)-mediated translation inhibition in zebrafish and mouse early embryos (*17*, *18*). Here we report a parallel and conserved transcriptional innate immune response to dsRNA and its regulatory mechanisms in mESCs.

## dsRNA activates an ISG response in mouse stem cells

To gain a global insight into the transcriptomic effects of dsRNA in ESCs, we performed RNA-seq analysis on dsRNA-transfected mESCs and control mouse embryonic fibroblasts (MEFs) (table S1). In mESCs, 3755 differentially expressed genes (DEGs) were identified (Fig. 1A). Gene Ontology (GO) enrichment analysis of top 100 upregulated dsRNA-responsive genes revealed significantly enriched terms, including “response to virus” and “cellular response to interferon-β (IFN-β)”, among others (Fig. 1B). Notably, a subset of interferon-stimulated genes (ISGs) was strongly induced, such as *Isg15*, *Cxcl10*, *Oasl1*, and *Ifih1* (Fig. 1C). In contrast to MEFs, no interferon (IFN) or IFN receptor genes were upregulated in mESCs upon dsRNA stimulation (table S1). We further observed that the dsRNA-induced ISG response in mESCs was dose-dependent (fig. S1A). Additionally, a panel of p53 target genes was activated, including *Id2*, *Mdm2*, *Gdf15*, *Pmaip1*, *Trp53inp1*, and *Bbc3*, indicating p53 activation. The dsRNA-mediated induction of ISGs and p53 target genes was validated by quantitative polymerase chain reaction (q-PCR) analyses (Fig. 1D). Importantly, the ISG response and p53 target activation were independent of *de novo* protein translation, as it remained unaffected by cycloheximide (CHX), a specific translation inhibitor (fig. S1B). Taken together, these results demonstrate that dsRNA stimulates ISG expression in mESCs through an IFN-independent pathway.

**Figure 1.**
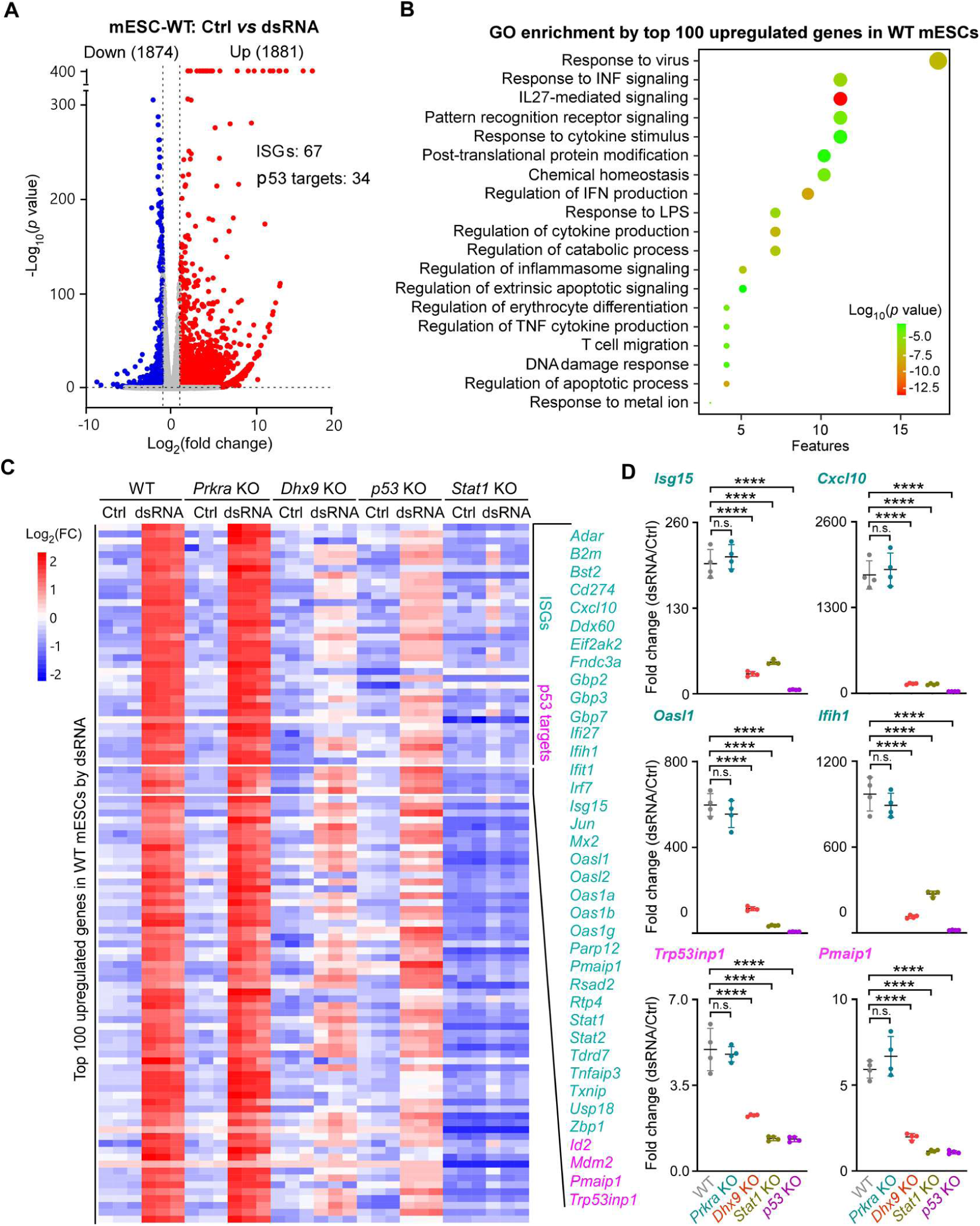
dsRNA activates an ISG response in mESCs. (**A**) Volcano plots of differentially expressed genes in control and dsRNA transfected mESCs. Dashed lines indicate fold change (log_2_FC > 1.0) and *p* value cutoffs (p < 0.05). WT, wild type. (**B**) GO enrichment analyses of the dsRNA induced top 100 genes identified in (A). WT, wild type. (**C**) Heatmap of the expression of the dsRNA induced top 100 genes in WT mESCs in control or dsRNA transfected *Prkra* KO, *Dhx9* KO, *p53* KO, and *Stat1* KO mESCs (n = 3 for each group). FC, fold change; WT, wild type; KO, knockout; Ctrl, control. (**D**) RT-qPCR results showing expressional fold changes of the indicated ISG genes (*Isg15*, *Cxcl10*, *Oasl1*, and *Ifih1*) and p53 target genes (*Pmaip1* and *Trp53inp1*) in dsRNA transfected *vs* control mESCs of WT, *Prkra* KO, *Dhx9* KO, and *p53* KO. Ordinary one-way ANOVA with multiple comparisons. n.s., not significant. ****, *p* < 0.0001. Ctrl, control; WT, wild type; KO, knockout.

## Dhx9 is a dsRNA stress sensor to activate ISGs expression

We previously identified Prkra as a dsRNA sensor that inhibits translation in zebrafish and mouse ESCs (*17*, *18*). However, dsRNA-induced ISG expression remained unaffected in *Prkra*-knockout (KO) ESCs (Fig. 1C and table S2), indicating the involvement of additional dsRNA sensor(s) in this signaling pathway. To identify such sensor(s), we performed a pull-down assay using the J2 dsRNA-specific antibody on the cytoplasmic fraction of mESCs at 12 hours post-dsRNA transfection, followed by mass spectrometry (MS) analysis for protein identification (Fig. 2A). As anticipated, several canonical dsRNA-binding proteins (dsRBPs) were enriched in the screen, including the RNA helicases Dhx9 and Ddx3x, the RNase Dicer, the dsRNA-specific adenosine deaminase Adar, as well as Prkra, Tarbp2, Tdp43, and Stau1 (Fig. 2A). The tumor suppressor p53 was also detected in the dsRNA-protein complex (Fig. 2A).

**Figure 2.**
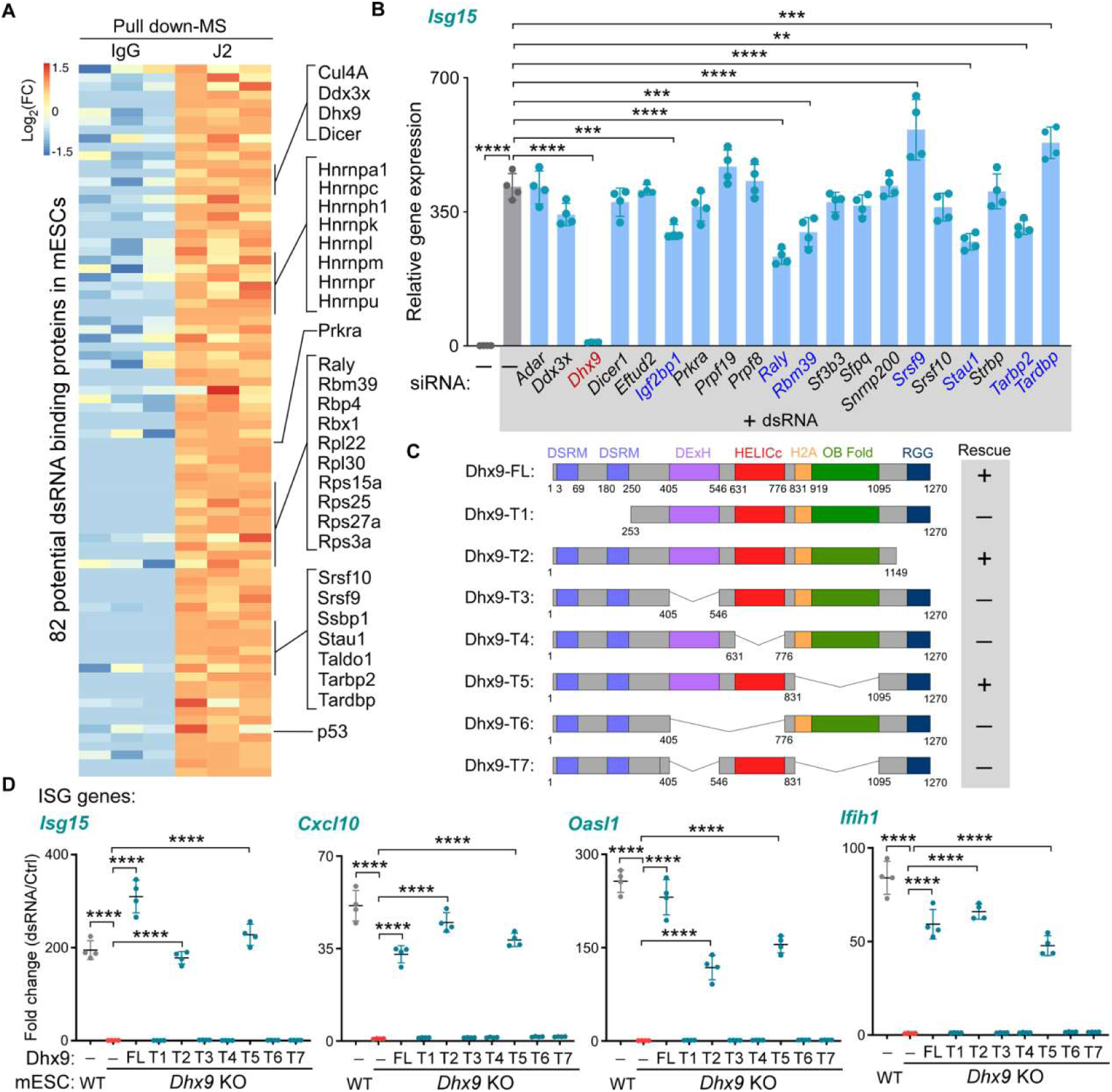
Dhx9 is a dsRNA stress sensor to activate ISGs expression in mESCs. (**A**) The proteins significantly enriched in the J2 antibody pulldown samples (n = 3) compared to control IgG samples (n = 3) with the statistical cut-offs of *p* value < 0.01 and fold change > 2. MS, mass spectrometry; FC, fold change. (**B**) RT-qPCR results showing the relative expression level of *Isg15* in control or dsRNA transfected mESCs when the indicated genes were knocked down by their siRNAs. Ordinary one-way ANOVA with multiple comparisons. **, *p* < 0.01; ***, *p* < 0.001; ****, *p* < 0.0001. (**C**) Schematic representation of the structures of mouse Dhx9 truncates used in the rescue assays. FL, full length; T, truncate. (**D**) RT-qPCR results showing expressional fold changes of the indicated ISG genes (*Isg15*, *Cxcl10*, *Oasl1*, and *Ifih1*) in dsRNA transfected *vs* control mESCs of WT or *Dhx9* KO with the indicated Dhx9 truncates overexpressed. Ordinary one-way ANOVA with multiple comparisons. n.s., not significant. ****, *p* < 0.0001. WT, wild type; KO, knockout; Ctrl, control.

We next investigated which of these dsRBPs functions as the sensor mediating dsRNA-induced ISG responses in mESCs. Specific siRNAs targeting the different dsRNA binding proteins were transfected into mESCs, following by dsRNA transfection and RT-qPCR analysis of the induction of the ISGs. The results showed that knockdown of Dhx9 dramatically inhibited the dsRNA induced ISG response, while knockdown of other dsRNA binding proteins tested has much weaker or no clear effects, implicating Dhx9 as the key sensor (Fig. 2B). The knockdown effects of the siRNAs on the target mRNA expression levels were verified by RT-qPCR (fig. S2). We then created a *Dhx9* knockout mESC line using the CRISPR/Cas9 system (fig. S3A) and confirmed that *Dhx9* mutation markedly suppresses the dsRNA-induced activation of either ISGs or p53 target genes (Fig. 1, C and D and table S2). Dhx9 harbors a conserved helicase core domain, two N-terminal dsRNA-binding domains (dsRBDs), an oligonucleotide/oligosaccharide-binding fold (OB-fold), a nuclear transport domain, and a C-terminal RGG-box motif with single-stranded DNA-binding activity (*19*). To define the functional domains of Dhx9 required for ISGs and p53 target genes activation, we generated a series of Dhx9 deletion constructs and assessed their ability to rescue dsRNA-induced ISG expression in *Dhx9* KO ESCs (Fig. 2C). Our results demonstrated that the dsRBDs and helicase domain are essential for ISGs and p53 target induction, whereas the OB-fold and RGG-box are dispensable (Fig. 2, C and D and fig. S3B).

## dsRNA stimulation stabilizes Stat1 and p53 to activate the ISGs

Given that numerous ISGs are direct transcriptional targets of Stat1 (*20*), we investigated whether Stat1 is activated in dsRNA-transfected mESCs. Intriguingly, dsRNA transfection led to a dramatic increase in total Stat1 protein levels, whereas the tyrosine 701 (Tyr701)-phosphorylated form of Stat1 remained unchanged (Fig. 3A). Western blotting analysis confirmed that dsRNA treatment also upregulated p53 protein levels in mESCs, further supporting p53 activation (Fig. 3A). Critically, dsRNA induced significantly lower ISG expression in mESCs lacking either *p53* or *Stat1* than in wild-type (Fig. 1, C and D; fig. S3A; and table S2), indicating that both molecules are essential for the dsRNA-mediated ISG response.

**Figure 3.**
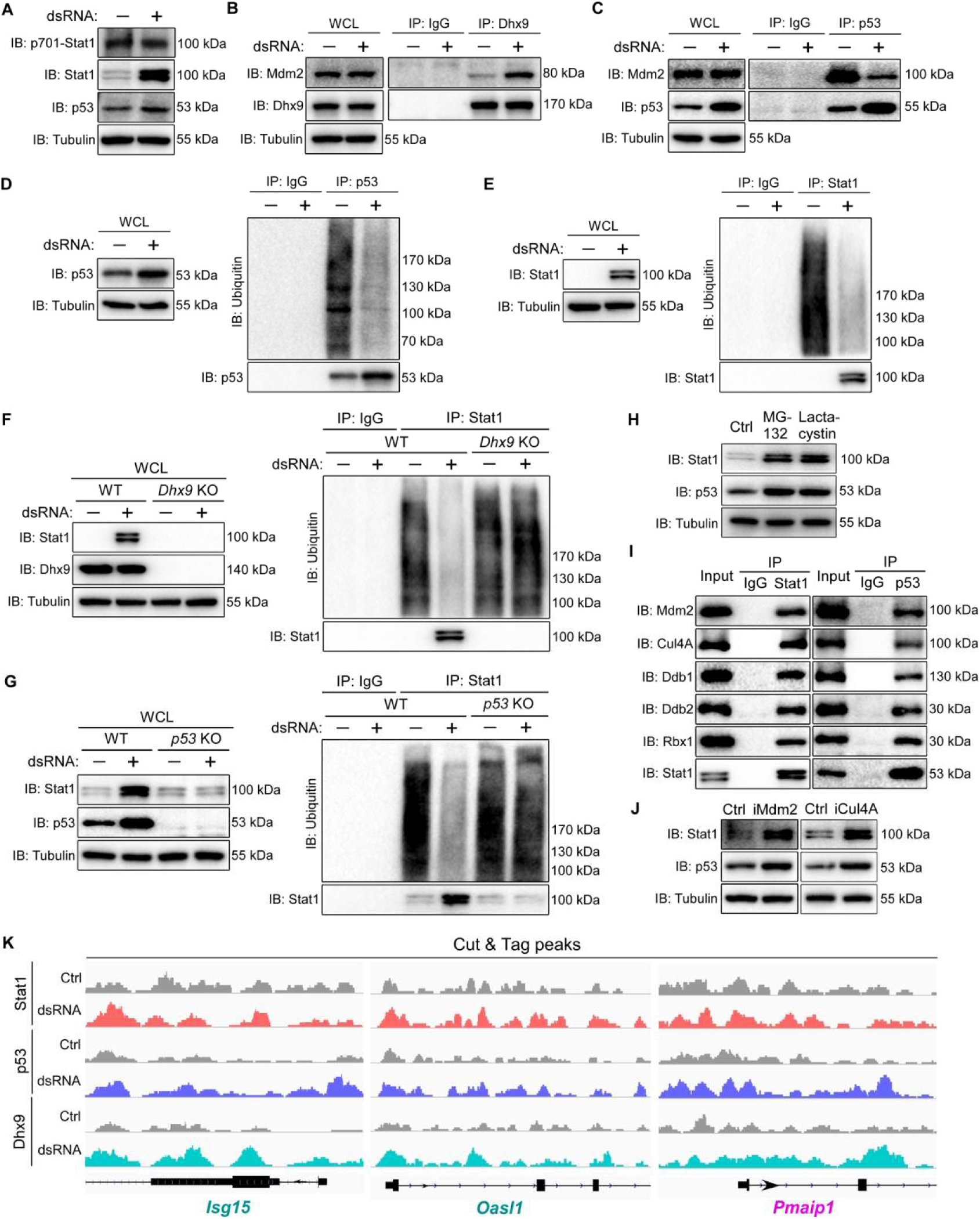
dsRNA stimulation stabilizes Stat1 and p53, which cooperate with Dhx9 to activate the ISGs in mESCs. (**A**) Immunoblotting images showing the expression of indicated protein in control or dsRNA transfected WT mESCs. IB, immunoblotting. (**B**, **C**) Co-IP assays showing the interactions of Dhx9-Mdm2 (B) or Mdm2-p53 (C) in control or dsRNA transfected WT mESCs. WCL, whole cell lysate; IP, immunoprecipitation; IB, immunoblotting. (**D**, **E**) *In vivo* ubiquitination analysis of p53 (D) or Stat1 (E) in control or dsRNA transfected WT mESCs. WCL, whole cell lysate; IP, immunoprecipitation; IB, immunoblotting. (**F**, **G**) *In vivo* ubiquitination analysis of Stat1 in control or dsRNA transfected mESCs of WT, *Dhx9* KO (F), or *p53* KO (G). WCL, whole cell lysate; IP, immunoprecipitation; IB, immunoblotting; WT, wild type; KO, knockout. (**H**) Immunoblotting images showing the expression of indicated protein in MG132 or Lactacystin treated WT mESCs. Ctrl, control; IB, immunoblotting. (**I**) Co-IP assays showing the interactions between Stat1 or p53 and Mdm2 or CRL ubiquitin ligase machinery. IP, immunoprecipitation; IB, immunoblotting. (**J**) Immunoblotting images showing the expression of indicated protein in Nutlin (Mdm2 inhibitor, iMdm2) or KH-4-43 (Cul4A inhibitor, iCul4A) treated WT mESCs. IB, immunoblotting. (**K**) Cut&Tag peaks (visualized by IGV) of Dhx9, p53, and Stat1 for the selected ISG genes (*Isg15* and *Oasl1*) or p53 target gene (*Pmaip1*) in control or dsRNA transfected WT mESCs. Ctrl, control.

We next explored the mechanism by which Dhx9 facilitates the stabilization of p53 and Stat1. p53 is highly expressed in ESCs and plays a key role in regulating self-renewal and pluripotency (*21-24*). In normal cells, p53 protein levels are maintained at a low baseline *via* ubiquitination and subsequent proteasomal degradation, a process mediated by E3 ubiquitin ligases such as Mdm2 (*25-28*). Notably, Mdm2 has been reported to interact with Dhx9, and this interaction can be further enhanced by long non-coding RNAs (lncRNAs) (*29*). We hypothesized that formation of the Dhx9-dsRNA complex disrupts the Mdm2-p53 interaction, thereby stabilizing and activating p53. To test this, we performed co-immunoprecipitation (co-IP) assays and found that dsRNA transfection significantly increased the amount of Mdm2 pulled down with Dhx9 (Fig. 3B), while concurrently reducing Mdm2-p53 binding (Fig. 3C) and p53 ubiquitination (Fig. 3D).

We then examined whether dsRNA-induced Stat1 stabilization in mESCs is regulated *via* the ubiquitin-proteasome pathway. Indeed, Stat1 underwent extensive ubiquitination and degradation in naive mESCs, whereas dsRNA transfection markedly reduced Stat1 ubiquitination and promoted its stabilization (Fig. 3E). Unexpectedly, both Dhx9 and p53 were required for this dsRNA-mediated Stat1 stabilization (Fig. 3, F and G). We reasoned that this process is post-translational—likely mediated by ubiquitination regulation, analogous to p53, and orchestrated by the Dhx9-dsRNA complex. Indeed, proteasome inhibitor treatment efficiently stabilized both p53 and Stat1 (Fig. 3H). To identify the E3 ubiquitin ligase responsible for Stat1 regulation, we revisited our proteomic analysis of dsRNA-associated factors in mESCs and found that components of the CRL4 E3 ligase complex—including Cul4A and its catalytic subunit Rbx1—were among the most highly enriched proteins (Fig. 2A). We confirmed the interaction between Stat1 and Cul4A in mESCs *via* co-IP analysis (Fig. 3I). Interestingly, Cul4A has been suggested to associate with Mdm2 to promote p53 proteolysis (*30*). We assumed that Cul4A and Mdm2 likely cooperate to degrade Stat1 and p53 in mESCs. Indeed, in mESCs treated with MG132, immunoprecipitation with either p53 or Stat1 efficiently pulled down both Mdm2 and Cul4A, together with other CRL4 E3 ligase components (Fig. 3I). Further, treatment of mESCs with specific inhibitors of either Mdm2 or Cul4A stabilized both p53 and Stat1, leading to ISGs and p53 target genes activation (Fig. 3J and fig, S4, A and B).

Collectively, these data suggest a model that both p53 and Stat1 are constitutively ubiquitinated by a Mdm2/Cul4A machinery and degraded in mESCs, and that dsRNA stimulation—dependent on Dhx9—largely blocks this degradation pathway to stabilize both proteins and activate the ISG response.

## Stat1, Dhx9 and p53 cooperate to activate the ISGs

Although Stat1 activity is maximized upon phosphorylation, unphosphorylated Stat1 (U-Stat1) can also translocate into the nucleus and regulate the expression of immune-related genes (*31*). In mouse macrophages, RNA virus infection can induce Dhx9-Stat1 interaction which then cooperatively promotes the transcription of ISGs (*32*). Building on these observations, we hypothesized that dsRNA-stabilized U-Stat1 might collaborate with Dhx9 and p53 to drive ISGs activation in mESCs. To test this, we mapped the chromatin occupancy of Stat1, Dhx9, and p53 in dsRNA-transfected mESCs using CUT&Tag (Cleavage Under Targets & Tagmentation) assays. Consistent with our hypothesis, the occupancy of all three proteins was significantly enhanced at the promoter regions of top dsRNA-induced ISGs and p53 target genes, with *Isg15*, *Oasl1*, and *Pmaip1* serving as representative examples (Fig. 3K). Interestingly, the three factors may occupy overlapping or different sites in the ISG genes (Fig. 3K and fig. S4C). Of note, Stat1 is also robustly transcriptionally induced by dsRNA, establishing a positive feedback loop that further amplifies the response (Fig. 1C).

## Dhx9 regulates the p53/Stat1 destruction machinery in the dsRNA induced condensates

The organization into liquid-like condensates through phase separation is a common feature for many signaling complexes in innate immunity, which is critical for the activation of downstream events (*33*). In mammalian cells, cytosolic dsRNA has been reported to trigger the formation of dsRNA-protein condensates named dsRNA-induced foci (dRIFs) through phase separation and dRIF formation has been suggested to be involved in PKR activation by dsRNA stress (*34*). We examined the subcellular localization of Dhx9 in dsRNA transfected cells. Interestingly, although Dhx9 localizes predominantly in the nuclear in control mESCs, they became abundant in the dRIFs where they co-localize well with dsRNAs in transfected cells (Fig. 4A). Subcellular fractionation experiment also showed that the cytoplasmic Dhx9 greatly increased upon dsRNA transfection (fig. S5A).

**Figure 4.**
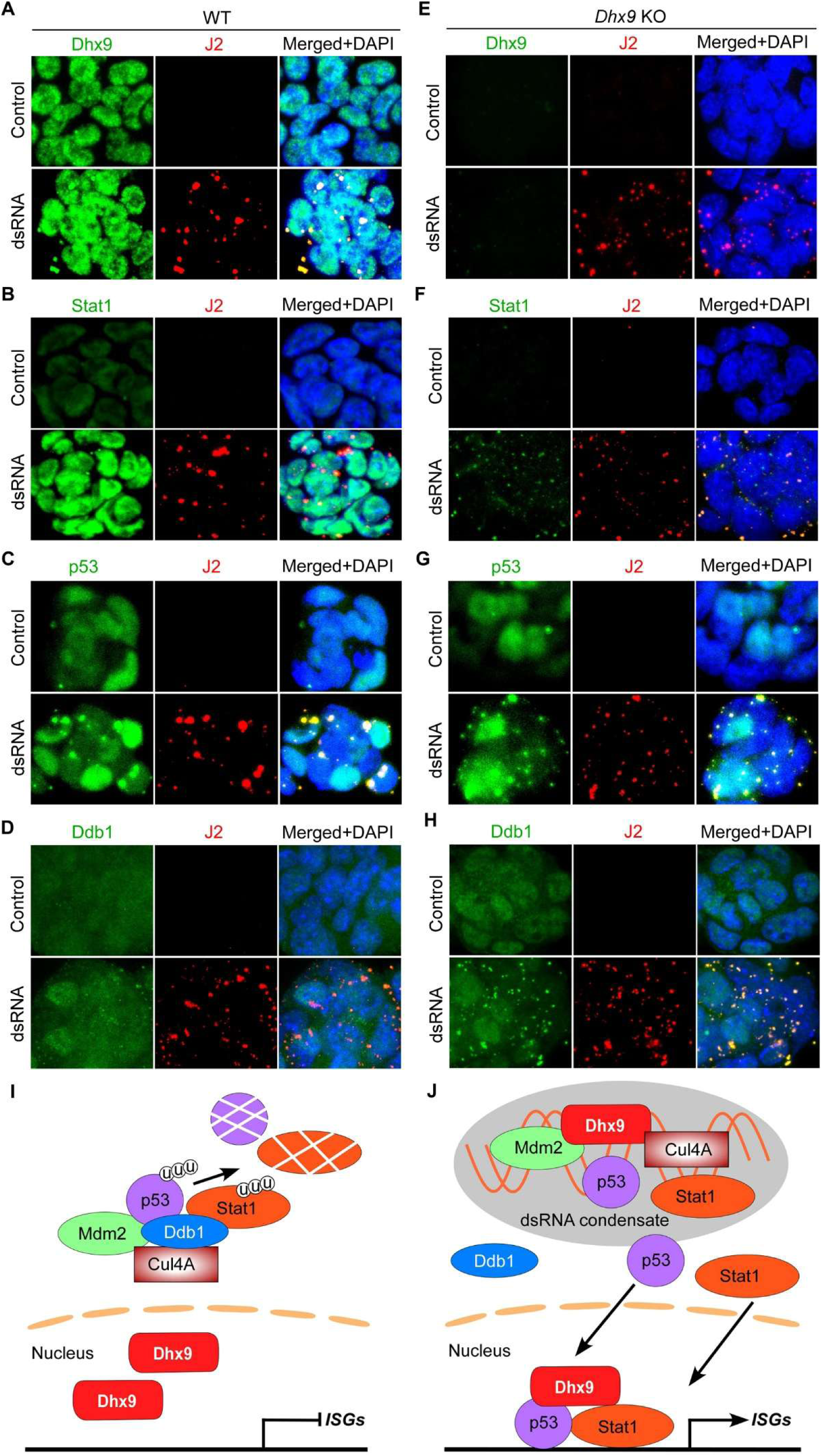
Dhx9 regulates the p53/Stat1 destruction machinery in the dsRNA induced condensates in mESCs. (**A-D**) Representative immunofluorescence images showing the co-staining of dsRNA (J2) and Dhx9 (A), Stat1 (B), p53 (C), or Ddb1 (D) in WT mESCs. WT, wild type. (**E-H**) Representative immunofluorescence images showing the dsRNA foci (E) and their co-staining with Stat1 (F), p53 (G), or Ddb1 (H) in Dhx9 KO mESCs. KO, knockout. (**I**, **J**) A model for the Dhx9-dsRNA mediated p53, Stat1 stabilization and ISG activation. In wild type mESCs, p53 and Stat1 are ubiquitinated and thus degradation by Mdm2/Cul4A containing E3 ligase complex (I). Dhx9 senses dsRNA to recruit the p53/Stat1 ubiquitination complex into the condensates while excluding their adaptor Ddb1, leading to release and stabilization of p53 and Stat1 (J).

We assume that the formation of the dRIFs is important for the stabilization of Stat1 and ISGs activation. Indeed, Dhx9, p53, mdm2, Stat1, Cul4A were all detected in the dRIFs (Fig. 4, A-C and fig. S5. B and C). In the absence of Dhx9, the dRIFs still formed, yet their sizes were smaller (Fig. 4). p53, Stat1 were also present in the dRIFs without Dhx9, but they were no longer stabilized as evidenced by the near-complete loss of their nuclear localization (Fig. 4, B, C, F and G). Thus, Dhx9 should be important for the control of size of the dRIFs and the p53/Stat1 ubiquitination reaction in the dRIFs. One possibility is that p53/Stat1 might be sequestered from the ubiquitin ligases in the dRIFs. By immunofluorescent staining, p53, Cul4A and Mdm2 were strongly enriched in the dRIFs in dsRNA transfected mESCs, either in the presence or absence of Dhx9 (Fig. 4, C and G; fig. S5, B-E). For Stat1, it is strongly detected in the dRIFs in *Dhx9* KO cells, and relatively weakly in wild type ESCs (Fig. 4, B and F). However, it is readily detected in the dsRNA immunoprecipitants by Western blotting analysis (fig. S6A). We then tested other components of the CRL4 ubiquitination complex in the dRIFs upon dsRNA treatment in wild type and *Dhx9* KO or *p53* KO mESCs. The results showed that damage-specific DNA binding protein 1 (Ddb1), a key substrate adapter protein for Cul4a (*35*), exhibited only mild enrichment in dRIFs following dsRNA stimulation in mESCs (Fig. 4D). Interestingly, in the *Dhx9* KO and *p53* KO mESCs, Ddb1 became efficiently recruited to the dRIFs, likely restored the CRL4 activity for Stat1 and p53 ubiquitination (Fig. 4H and fig. S5F). Biochemically, the Cul4A associated Ddb1 is much reduced in the dsRNA transfected cells compared with untreated or *Dhx9* KO and *p53* KO cells, where Stat1 ubiquitination was unaffected (fig. S6B). Depletion of Ddb1 has been reported to stabilize p53 in ESCs (*36*). These data suggest a model that the dRIFs recruit the p53/Stat1 ubiquitination complex into the condensates while excluding their adaptor Ddb1, leading to release and stabilization of p53 and Stat1 (Fig. 4, I and J).

## The dsRNA-Dhx9-p53-Stat1 cascade functions in stem cell under dsRNA stress

Virus is the most important source of exogenous dsRNA to stimulate the innate immune response in differentiated cells (*37*). In tissue stem cells, the RNAi pathway has been shown to be important for defense against RNA virus infection (*38*), including Zika virus (ZIKV), a single-stranded RNA virus of the Flaviviridae family that would produce dsRNA intermediates during replication (*39*). We tested whether the Dhx9 mediated ISG response functions in antiviral defense in mESCs against ZIKV. We demonstrated that ZIKV is capable of infecting mESCs, undergoing efficient replication and resulting in dsRNA accumulation within mESCs (Fig. 5A). Notably, ZIKV infection triggered the expression of ISGs and p53 target genes in a Mavs-independent manner in mESCs, whereas this inductive effect was completely abolished in *Dhx9* KO mESCs (Fig. 5B). Importantly, loss of Dhx9 but not Mavs dramatically enhanced ZIKV RNA replication in mESCs (Fig. 5, A, C and D), supporting a crucial role of Dhx9-mediated immunity in virus defense in mESCs.

**Figure 5.**
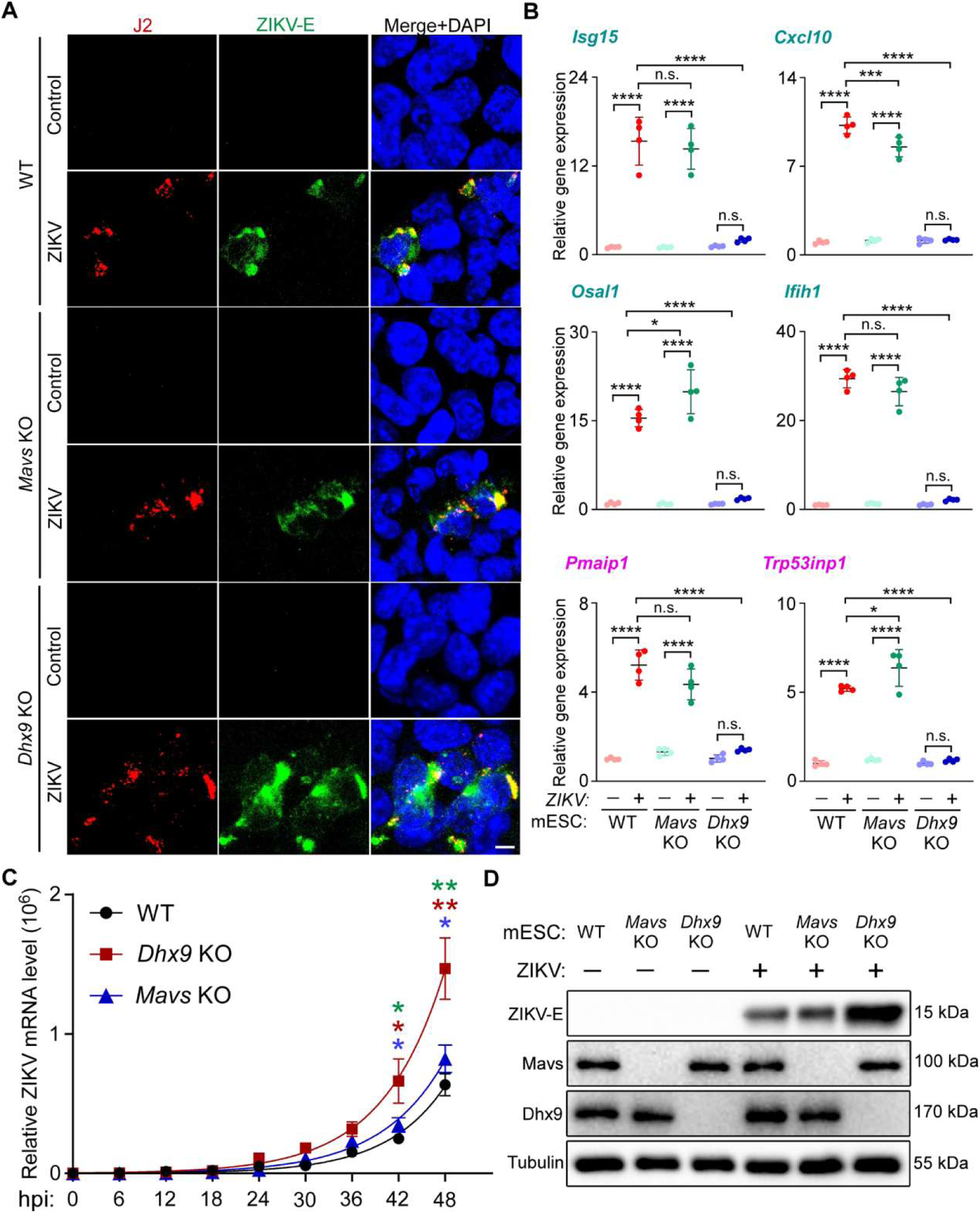
Dhx9 mediates Mavs-independent innate defense against Zika virus in mESCs. (**A**) Representative immunofluorescence images showing dsRNA (J2) and ZIKV envelope protein (ZIKV-E) in WT, *Mavs* KO and *Dhx9* KO mESCs at 48 hpi. hpi, hours post infection; WT, wild type; KO, knockout. Scale bar, 5 μm. (**B**) RT-qPCR results showing the relative expression level of ISGs (*Isg15*, *Cxcl10, Oasl1, and Ifit8*) and p53 targets (*Pmaip1* and *Trp53inp1*) in WT or *Dhx9* KO mESCs infected by ZIKV at MOI of 0.5 and 1.0 at 48 hpi. Ordinary one-way ANOVA with multiple comparisons. n.s., not significant. *, *p* < 0.05; ***, *p* < 0.001; ****, *p* < 0.0001. WT, wild type; KO, knockout. (**C**) RT-qPCR results showing the relative ZIKV mRNA level in WT, *Dhx9* KO, and *Mavs* KO mESCs at the indicated time point after infection. hpi, hours post infection; WT, wild type; KO, knockout. Students’*t* test. *, *p* < 0.05; **, *p* < 0.01. Significant differences between WT and *Dhx9* KO were labeled in red, between WT and *Mavs* KO in blue, and between *Dhx9* KO and *Mavs* KO in green. (**D**) Western blotting results showing the expression level of ZIKV envelope protein in WT, *Mavs* KO, and *Dhx9* KO mESCs infected by ZIKV at MOI of 1.0 at 48 hpi. hpi, hours post infection; WT, wild type; KO, knockout.

Mitochondrion is a common source of endogenous dsRNAs due to the bi-directional transcription of the mtDNA. Under stress conditions, mitochondrial (mt) dsRNAs may accumulate and be released into the cytoplasm, thereby triggering an innate immune response (*40*, *41*). We tested whether knockdown of *Pnpt1*, which encodes PNPase that is crucial in mt-dsRNA turnover (*42*), would activate the Dhx9 dependent ISG response in mESCs. Indeed, knockdown of *Pnpt1* by two independent shRNAs both stimulated the expression of a panels of ISGs and p53 target genes, including *Isg15*, *Cxcl10*, *Oasl1* and *Ifih1*, but not in *Dhx9* KO mESCs (fig. S7A).

Inhibition of monopolar spindle 1 (MPS1)—a master regulator of the spindle assembly checkpoint—has been demonstrated to trigger chromosome missegregation and the accumulation of cytoplasmic dsRNA derived from non-exonic transcripts (*43*). We thus investigated whether treatment with BAY-1217389, a selective MPS1 inhibitor, would elicit dsRNA accumulation and the activation of ISGs and p53 target genes in mESCs. Our results showed that BAY-1217389 treatment robustly induced an ISG response and p53 target activation in wild-type mESCs (fig. S7B). Notably, these effects were completely abrogated in *Dhx9* KO mESCs, providing compelling evidence that Dhx9 is required for sensing endogenous dsRNA in this context (fig. S7B).

Collectively, these results establish that Dhx9 is indispensable for ISG induction in mESCs following viral or endogenous dsRNA stimulation, thereby mediating antiviral and dsRNA stress responses.

## Evolution of the Dhx9-mediated dsRNA signaling cascade

We tested whether the Dhx9-mediated dsRNA sensing cascade is conserved in human ESCs. The results showed that dsRNA transfection also stimulates the expression of ISGs in hESCs, including *ISG15*, *CXCL10*, *IFIH1*, and *OASL* (Fig. 6, A-D and table S3). And DHX9, TP53 and STAT1 were also all observed in the dRIFs in these transfected cells (Fig. 5E and fig. S8, A and B). As in mESCs, dsRNA transfection or treatment with Mdm2/Cul4A inhibitors also stabilized p53 and STAT1, as well as ISGs and p53 target genes activation in hESCs (Fig. 5, F and G; fig. S8, C and D).

**Figure 6.**
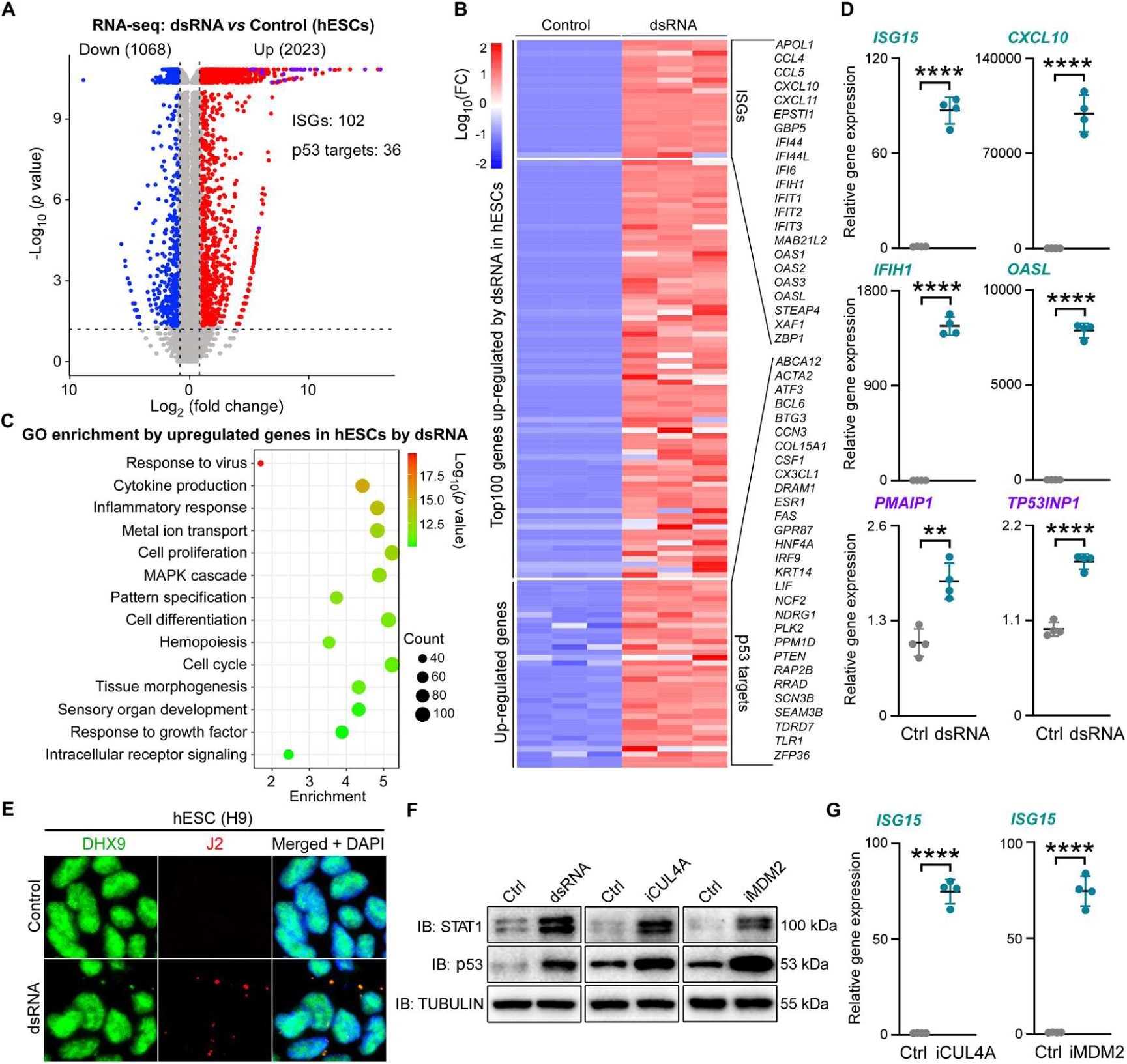
hESCs exhibit a similar ISG response to dsRNA via DHX9, p53, and STAT1. (**A**) Volcano plots of differentially expressed genes in control and dsRNA transfected hESCs. Dashed lines indicate fold change (log_2_FC > 1.0) and *p* value cutoffs (p < 0.05). (**B**) Heatmap of the expression of the dsRNA induced top 100 genes and p53 target genes in control (n = 3) or dsRNA (n = 3) transfected hESCs. FC, fold change. (**C**) GO enrichment analyses of the dsRNA induced top 100 genes identified in (A and B). (**D**) RT-qPCR results showing relative expression levels of the indicated ISGs (*ISG15*, *CXCL10*, *OASL1*, and *IFIH1*) and p53 target genes (*PMAIP1* and *TRP53INP1*) in control and dsRNA transfected hESCs. Student’s *t*-test. **, *p* < 0.01; ****, *p* < 0.0001. Ctrl, control. (**E**) Representative immunofluorescence images showing the co-staining of dsRNA (J2) and DHX9 in hESCs. (**F**) Immunoblotting images showing the expression of indicated proteins in Nutlin (MDM2 inhibitor, iMDM2) or KH-4-43 (CUL4A inhibitor, iCUL4A) treated hESCs. IB, immunoblotting. (**G**) RT-qPCR results showing relative expression levels of *ISG15* in Nutlin (MDM2 inhibitor, iMDM2) or KH-4-43 (CUL4A inhibitor, iCUL4A) treated hESCs. Student’s *t*-test. ****, *p* < 0.0001. Ctrl, control.

In differentiated Neuro-2a (N2A) cells, knockdown of either *Dhx9* or *Mavs* reduced dsRNA-triggered ISGs induction, including *Isg15*, *Cxcl10*, *Ifih1*, and *Oasl1* (fig. S8E). Furthermore, double knockdown of *Dhx9* and *Mavs* elicited a much stronger effect, suggesting that the dsRNA-Dhx9 cascade likely also contributes to the dsRNA response in differentiated cells. We tested whether the above mechanism is conserved in zebrafish embryos. We analyzed RNA-seq data of 6 hpf embryos injected with dsRNAs and identified 741 upregulated and 985 downregulated genes in dsRNA-injected samples (Fig. 7, A and B; table S4). Several core downstream genes of the p53 pathway, including *phlda3* and *cdkn1a* (*p21*), were upregulated as confirmed by RT-qPCR, suggesting activation of the p53 pathway (Fig. 7, B and C). In addition, 24 ISGs but no IFN ligand genes were also stimulated (Fig. 7B and table S4), indicating ISG gene upregulation independent of IFN signaling. We confirmed the explosive upregulation of the ISGs, including *isg15*, *casp8*, *cxcl12*, and *ifit8*, in dsRNA injected embryos by RT-qPCR (Fig. 7C) but failed to observe any increased expression of all the five IFN ligand genes in zebrafish (fig. S9A). The dsRNA induced ISGs stimulation is directly controlled by a maternal mechanism, as CHX treatment from 2 hpf to 6 hpf, which blocks protein synthesis after zygotic genome activation, could not prevent the induction of the ISGs and p53 target, such as *isg15, casp8*, *phlda2*, *ifit8*, *cdkn1a*, and *pdlha3*, except for *cxcl12b* (fig. S9, B-D).

**Figure 7.**
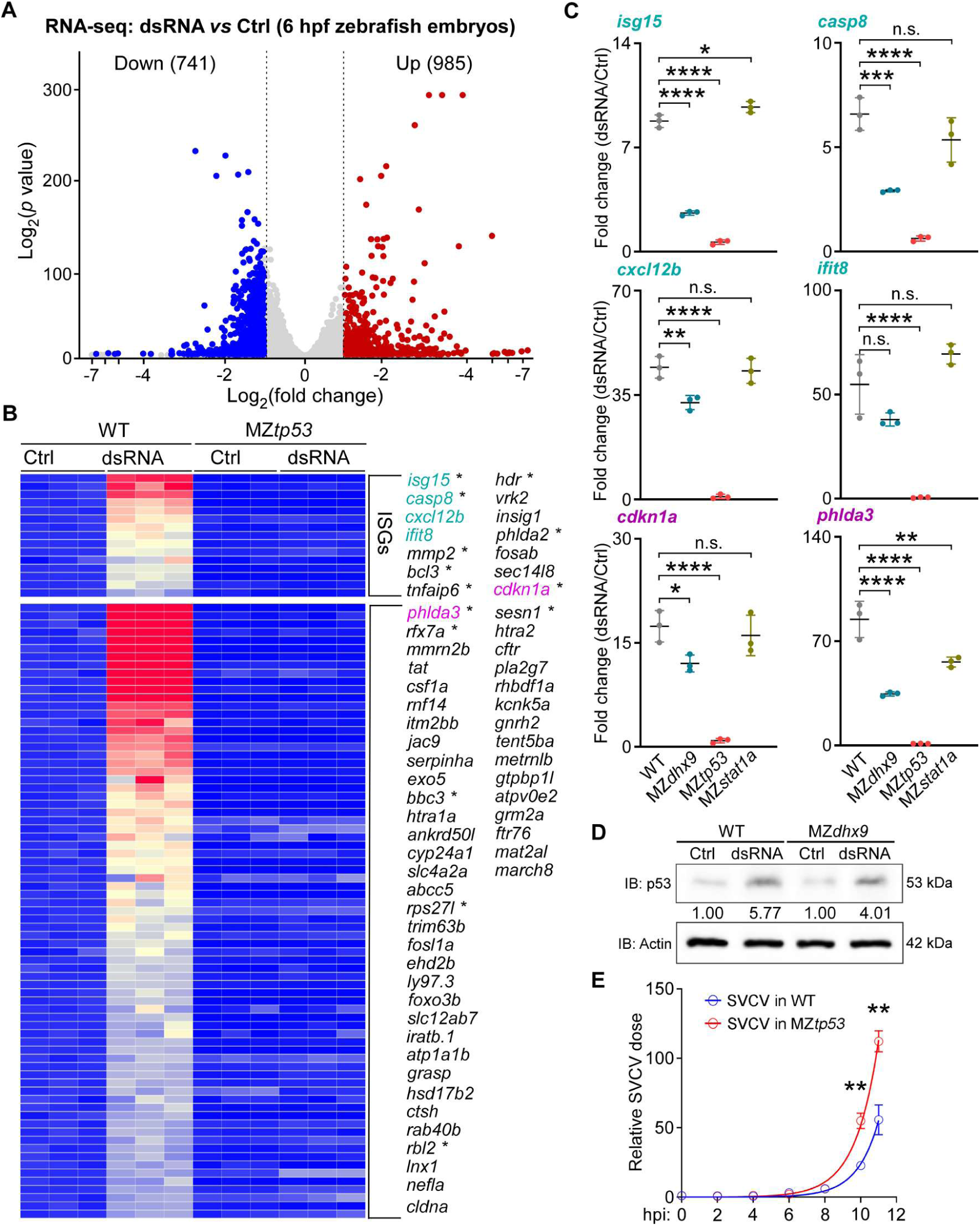
dsRNA induced an ISG response partially dependent on dhx9 in zebrafish early embryos. (**A**) Volcano plots of differentially expressed genes in control and dsRNA injected zebrafish embryos at 6 hpf. Dashed lines indicate fold change (log_2_FC > 1.0). Ctrl, control. hpf, hour post fertilization. (**B**) Heatmap of the expression of the dsRNA induced ISGs and other genes in control and dsRNA injected zebrafish embryos of WT or *tp53* mutant at 6 hpf (n = 3 for each group). Known p53 target genes were labeled with asterisks (*). WT, wild type; MZ*tp53*, maternal and zygotic *tp53* mutant; Ctrl, control. (**C**) RT-qPCR results showing expressional fold changes of the indicated ISG genes (*isg15*, *cxcl12b*, *casp8*, and *ifit8*) and p53 target genes (*phlda3* and *cdkn1a*) in dsRNA injected *vs* control zebrafish embryos of WT, *dhx9* KO, *stat1a* KO and *tp53* mutant. Ordinary one-way ANOVA with multiple comparisons. n.s., not significant. ****, *p* < 0.0001. Ctrl, control; WT, wild type; MZ*tp53*, maternal and zygotic *tp53* mutant; MZ*dhx9*, maternal and zygotic *dhx9* knockout; MZ*stat1a*, maternal and zygotic *stat1a* knockout. (**D**) Immunoblotting images showing the expression level of p53 protein in control or dsRNA injected zebrafish embryos of WT and *dhx9* KO at 6 hpf. IB, immunoblotting; Ctrl, control; WT, wild type; MZ*dhx9*, maternal and zygotic *dhx9* knockout. (**E**) Relative SVCV dose in WT and MZ*tp53* zebrafish embryos at the indicated time points after SVCV injection. WT, wild type; MZ*tp53*, maternal and zygotic *tp53* mutant. Students’s *t* test. **, *p* < 0.01.

As in mESCs, such a transcriptional response to dsRNA was also dependent on maternally supplied p53 in zebrafish embryos. We confirmed that p53 protein increased dramatically after injecting *cas9* dsRNA at 4 hpf, (Fig. 7D). Of the 741 genes induced by dsRNA in wild-type embryos, 616 (71.8%) failed to be upregulated in the maternal-zygotic *tp53* mutant (MZ*tp53*) background (table S4), including the canonical p53 target genes such as *phlda3, cdkn1a*, and fifteen ISGs (Fig. 7B and table S4). The results were confirmed by RT-qPCR (Fig. 7C). On the other hand, in *dhx9* KO zebrafish embryos (MZ*dhx9*), the dsRNA stimulated p53 activation and ISGs stimulation was only partially affected (Fig. 7C and fig. S10). The stabilization of p53 by dsRNA was also reduced in *dhx9* KO zebrafish embryos (Fig. 7, D). As Dhx9 was also present in the dsRNA immunoprecipitants in zebrafish embryos (*17*), we suggest Dhx9 likely is partially involved in the dsRNA response in zebrafish. There are two *stat1* orthologs in zebrafish, *stat1a* and *stat1b*. As Stat1b lacks a complete C-terminal transcription activation domain and is not expressed maternally (*44*), we tested whether Stat1a is involved in the dsRNA response in zebrafish embryos. In *stat1a* KO zebrafish embryos (MZ*stat1a*), the transcriptional effects of dsRNA on the ISGs remained unaffected, suggesting that Stat1 is not required for this process (Fig. 7C). Additionally, mutation of *tp53* significantly enhanced replication of spring viremia of carp virus (SVCV) in zebrafish embryos (Fig. 7E), implying an p53 dependent antiviral stress response in early zebrafish embryos.

## Discussion

In summary, we uncover a previously unrecognized dsRNA-sensing mechanism that drives an ISG response in ESCs. Cytoplasmic dsRNA acts as a molecular scaffold to recruit Dhx9 and other dsRNA-binding proteins, thereby triggering LLPS and the formation of cytoplasmic condensates. p53 and Stat1, together with their cognate ubiquitination machinery, are also recruited into these condensates. Owing to the remodeling of local molecular interactions within the condensates, Ddb1—a key adaptor protein for the ubiquitination of p53 and Stat1—is excluded from the complex. Consequently, p53 and Stat1 are stabilized, released from the condensates, and translocated to activate ISGs and p53 target genes transcription (Fig. 4, I and J). Because this pathway operates directly without additional signaling intermediates, it likely functions as an immediate, rapid-response mechanism to dsRNA stress.

In addition to its central role in gene regulation and RNA metabolism, Dhx9 has also been reported to function as a viral dsRNA sensor that mediates the IFN response in myeloid dendritic cells *via* MAVS (*45*). Conversely, Dhx9 can repress dsRNA-sensing pathways, likely by suppressing dsRNA accumulation or sequestering endogenous dsRNAs (*46*). In human cancer cells, upon infection with myxoma virus (MYXV), a member of the double-stranded DNA poxvirus family, DHX9 is recruited to cytoplasmic viral replication factories, forms unique granular cytoplasmic structures, and colocalizes with dsRNA to inhibit viral protein synthesis (*47*). More recently, Dhx9 has been shown to be recruited to cytoplasmic stress granules in response to UV-induced RNA damage, where it alleviates dsRNA stress (*48*, *49*). Our current work uncovers a unique role of Dhx9 in regulating p53/Stat1 signaling upon dsRNA stress in ESCs. We demonstrate that in ESCs, Mdm2 and components of the Cul4A E3 ligase form a ubiquitination complex responsible for the ubiquitination of p53 and Stat1. Consistent with this, CUL4A has been reported to be involved in MDM2-mediated proteolysis of p53 in human cell lines (*30*). Upon dsRNA stimulation, Dhx9 and the Mdm2/Cul4A complex are recruited to dsRNA foci, where Dhx9 is required for the exclusion of Ddb1 from the p53/Stat1 ubiquitination complex. Unexpectedly, p53 is also essential for the efficient exclusion of Ddb1 from the p53-Mdm2/Cul4A complex. The detailed changes in molecular interactions within the ubiquitination complex in the presence of dsRNA remain to be further elucidated.

In ESCs, the endogenous dsRNA level has been reported to be relatively high and the type I interferon response is inactive, due to lack of the classical dsRNA sensors, which is important for maintaining of the proliferation and pluripotency of stem cells (*50*). Here we show that upon high dose dsRNA stimulation, ESCs initiate an immediate p53/Stat1 dependent innate response. Such a pathway is responsive to high endogenous or viral dsRNAs and is crucial for virus defense in mESCs. This signaling pathway is conserved in human ESCs and likely also works in differentiated mammalian cells. In zebrafish embryos, p53 appears to be the major mediator of the transcriptional response to dsRNA, whereas Dhx9 contributes to a lesser extent, suggesting other dsRNA sensors may also be involved in this process.

We previously showed that Prkra mediates dsRNA-induced translational inhibition in zebrafish and mouse early embryos (*17*, *18*). Interestingly, Prkra is also recruited into the dsRNA condensates (Fig.2, A), which we suggest to be important for efficient translational inhibition. Thus, the dsRNA condensates formed by LLPS likely serve as a signaling center of the defensive dsRNA response in ESCs.

## Materials and methods

### Ethics statement

The experiments with infectious Zika virus (ZIKV) were conducted under biosafety level 2 (BSL2) facilities and approved by the Committee of Biosafety of Kunming Institute of Zoology, Chinese Academy of Sciences.

### Zebrafish, mRNA synthesis and microinjection

Zebrafish AB strain, tp53 mutant lines (*51*), as well as *dhx9* and *stat1a* CRISPants were maintained in a standard zebrafish breeding system (Haisheng) at 28 °C. All experiments were performed following the Animal Research: Reporting of *In Vivo* Experiments (ARRIVE) guidelines approved by the Ethics Committee for Animal Research of Life Science of Shandong University (permit number SYDWLL-2026-048).

Coding sequences of the genes of interest were PCR amplified or synthesized (TsingKe) and subcloned into the pCS2 vector. Recombinant plasmids were linearized with NotI, purified using a PCR cleanup kit, and used as templates for in vitro transcription. mRNA was synthesized in reactions containing cap analog (0.5 mM), ATP/CTP/UTP (0.5 mM each), GTP (0.1 mM), SP6 RNA polymerase (20 U), RNase inhibitor (20 U), 1× RNA polymerase buffer, and linearized DNA template, followed by incubation at 37 °C for 1.5 h. DNA templates were removed by DNase I digestion, and RNA was purified by phenol–chloroform extraction and isopropanol precipitation. The RNA pellet was washed with 70% ethanol and dissolved in RNase-free water.

Injection needles were prepared using a Narishige needle puller, and the tips were trimmed with fine tweezers to an appropriate diameter. Needles were mounted on a PLI-100A Picoliter Microinjector. IVT mRNAs, dsRNAs, or other reagents were loaded using the injector “Fill” function and injected into one-cell stage zebrafish embryos at a volume of 1-2 nL *per* embryo.

### *In situ* hybridization

ISH was performed as previously described (*52*). Probes for sox17, isg15 and stat1a were reported previously (*53*, *54*). Fragments of *sox17*, *isg15* and *stat1a* were amplified by PCR and cloned in pZeroback vector. Digoxigenin-labelled RNA probes were synthesized by *in vitro* transcription using DIG-labeling mix and SP6 or T7 RNA polymerase. Following DNase I treatment (Roche, 4716728001), probes were directly precipitated with 50% (v/v) isopropanol without phenol–chloroform extraction.

### Maternal and Zygotic gene knockout in zebrafish

We adopted the CRISPant strategy to rapidly generate zygotic mutant and maternal mutant (*55*), sgRNAs were designed by the online tools CRISPRscan. To generate dhx9 and stat1a zygotic mutant, the six high-efficient isg15 sgRNAs were equally mixed (50 ng/μL for each) and coinject with Cas9 protein (working concentration: 100 ng/μL) into the 1-cell stage embryos (2 nL for each). The adult CRISPant female F0 were crossed with the CRISPant male F0 to obtain Maternal-Zygotic mutant embryos. They were verified by RT-PCR analysis using the cDNAs from shield-stage embryos (maternal and zygotic transcripts analysis). The PCR products were subcloned and sent for sanger sequencing. The following sgRNAs for *dhx9* and *stat1a* were used: *dhx9*-sg1: 5’-GGGGAGAGTTACAGTTACAT-3’; *dhx9*-sg2: 5’- GGGGCAGTGGACAGTTAGAC-3’; *dhx9*-sg3: 5’- GGGACGTAATAATGCACCAT-3’; *dhx9*-sg4: 5’- GGTTGTGGTCAGGGCCCACT-3’; *dhx9*-sg5: 5’- GGGGTTACTGCTCGTGAGCA-3’; *dhx9*-sg6: 5’-GGCCGTCAGCTGTACCATTT-3’; *stat1a*-sg1: 5’- GGATATTGTCGGATGGCCAT-3’; *stat1a*-sg2: 5’- GGTTGTCTTGGAAGAGGTTC-3’; *stat1a*-sg3: 5’-GGTTTGCTCAGCGTCCTGAA-3’; *stat1a*-sg4: 5’- GGCCGAGGTGTTGAACCTGG-3’; *stat1a*-sg5: 5’-GGCAGTTGGGTCGGCATGCA-3’; and *stat1a*-sg6: 5’-GGGGATGGGCGTCCATCATG-3’. Primers used for genotyping for *tp53* mutants, as well as *dhx9* and *stat1a* CRISPants were: *dhx9*: forward, 5’- GAACTATGACATCCGAGCAG-3’ and reverse, 5’- ACAGCTGGTGTTGGACATCA-3’; *stat1a*: forward, 5’-TCAGCTACAGGAAGTGCTGA-3’ and reverse, 5’- GTGCTAATGTATCCCTCGTC-3’; and *tp53*: forward, 5’- ATTGCCAGAGTATGTGTCTGTCC-3’ and reverse, 5’-CTTAAAGGTGATGTCGGGTCT-3’.

### Cell lines and transfection

Mouse embryonic stem cells (mESCs) were used as our previously described (*17*). Briefly, mESCs were cultured on inactivated mouse embryonic fibroblast (MEF) feeder layers or 0.1% Gelatin (07903, Stemcell) in 2i medium composed of DMEM (C1199500BT, Gibco) supplemented with 15% FBS (A5669701, Gibco), 1× GlutaMAX (35050-061, Gibco), 1× non-essential amino acids (NEAA) (11140050, Gibco), 1× sodium pyruvate (11360-070, Gibco), 50 mM 2-mercaptoethanol (21985-023, Gibco), 1× penicillin-streptomycin (15140163, Gibco) and 10 ng/mL mouse leukemia inhibitory factor (mLIF) (ESG1107, Millipore), 3 μM CHIR99021 (S1263, Selleck) and 1 uM PD0325901 (S1036, Selleck). sgRNAs targeting the *Dhx9*, *Stat1*, or *p53* locus were introduced into mESCs by PX458 vector, and GFP-positive cells were sorted into 96-well plates by BD FacsAriaII Cell Sorters. Single clonal knockout cells were screened and confirmed by Sanger sequencing and expressions of target genes were examined by qRT-PCR and western blot analyses.

Human embryonic stem cell (hESC) line H9 was kindly provided by the Professor Bo Zhao (Kunming Institute of Zoology, Chinese Academy of Sciences) and were cultured with the Essential 8 hESC medium TeSR-E8 (05991, Stemcell) in Matrigel Matrix (354277, Corning) coated plates at 37 °C, 5% CO_2_, conditions. The medium was changed daily.

Mouse neuroblastoma N2a cells were cultured in Dulbecco’s Modified Eagle’s Medium (DMEM) (11965092, ThermoFisher Scientific) supplemented with 10% FBS (A5256701, ThermoFisher Scientific), 100 U/mL penicillin and 100 mg/mL streptomycin (A5873601, ThermoFisher Scientific). sgRNA-expressing cassettes targeting the *Mavs* and *Dhx9* genes were cloned into the pX458 vector, which expresses Cas9 ubiquitously. sgRNA vectors for each gene were transfected into the cells independently or together. Forty-eight hours post-transfection, dsRNA was introduced into the culture. After an additional 12 hours, the cells were harvested for total RNA extraction and qRT-PCR analysis.

HEK293T cells were cultured in Dulbecco’s Modified Eagle’s Medium (DMEM) (11965092, ThermoFisher Scientific) supplemented with 10% FBS (A5256701, ThermoFisher Scientific), 100 U/mL penicillin and 100 mg/mL streptomycin (A5873601, ThermoFisher Scientific).

Vero and C6/36 cells were used to virus propagation and titer determination. Vero cells (African green monkey kidney cells) were cultured in MEM (11095080, ThermoFisher Scientific) with 5% FBS (A5256701, ThermoFisher Scientific) at 37 °C. C6/36 cells (Aedes albopictus cells) were cultured in RPMI 1640 (11875093, ThermoFisher Scientific) with 10% FBS (A5256701, ThermoFisher Scientific) at 28 °C.

Lipofectamine 3000 (L3000015, ThermoFisher Scientific) were used for dsRNA transfections in ESCs and N2a cells while Lipofectamine 2000 (11668019, ThermoFisher Scientific) was used for siRNA transfection in ESCs following their manufacturer’s instructions respectively. Lipofectamine 2000 was used for plasmids transfection in HEK293T cells and a mouse embryonic stem cell nucleofector kit (VPH-1001, Lonza) was used to deliver plasmids into mouse ESCs.

### Virus propagation and infection

The Zika virus (ZIKV) strain MR766 (GenBank: LC002520) was kindly donated by Professor Cheng Tong from Xiamen University. Virus growth and titration was carried out as previously described (*56*). Briefly, the virus was propagated in C6/36 cells and titer was determined by a plaque assay on Vero cell. 80% confluent mESCs cultured in 24-well plate were infected by ZIKV at the MOI of 0.5, 1 or 5 for 6 hours followed by their complete growth medium. 48 hours after virus infection, mESCs were harvested for RT-qPCR assays. SVCV preparation and zebrafish embryo injection were used as our previously described (17).

### Plasmids and reagents

pX458 vector was used to CRISPR/Cas9 mediated genome editing to knockout target genes. The plasmid encoding full length human Dhx9 protein (Dhx9 FL) was a gift from professor Yuan Hu’s lab (Chongqing medical university). Various Dhx9 truncates were subcloned into pCMV-N-Flag vectors by PCR using the Dhx9 FL plasmid as templates. The obtained constructs were confirmed by Sanger sequencing and western blotting.

The following chemicals were used in this study. 1.5 μM MG132 (HY-13259, MedChemExpress) or 10 μM Lactacystin (HY-16594, MedChemExpress) was used to inhibit the proteasome-mediated protein degradation. 25 μM Nutlin-3a (HY-10029, MedChemExpress) or 50 μM KH-4-43 (HY-148602, MedChemExpress) was used to inhibit the activity of endogenous Mdm2 or Cul4a respectively. 10 to 1000 μM BAY1217389 (HY-12859, Selleck chemicals) was used to induce endogenous dsRNA accumulation. 10 μg/mL cycloheximide (CHX) (HY-12320, MedChemExpress) was used to inhibit translation in mESCs. poly(I:C) (tlrp-pic, InvivoGen) was used as a dsRNA mimic. Mouse or human ESCs were treated with each of above chemicals for 12 hours before being harvested for further analysis.

### Lentivirus preparation and infection

Lentivirus preparation and cell infection were carried out as previously described (*57*). The pLKO.1 lentiviral vectors were then co-transfected into HEK293T cells with the packaging plasmids pCMVΔ8.9 and pMD2.G at a ratio of 10:5:2. 48 hours after transfection, the culture medium was collected and centrifuged at 2500 rpm (3000 × *g*) for 10 min, and the supernatant was filtered. The supernatant containing lentiviral particles was centrifuged at 25,000 rpm (200,000 × *g*) at 4°C for 2.5 hours. The lentivirus pellet was then re-suspended in PBS containing 0.1% bovine serum albumin (BSA) and stored at -80°C in aliquots. The copy number of the lentiviral particles was confirmed *via* quantitative RT-PCR using the U5 primers (forward: 5’-AGCTTGCCTTGAGTGCTTCA-3’ and reverse: 5’-TGACTAAAAGGGTCTGAGGG-3’).

Mouse ESCs (5×10^5^) seeded in 6-cm dishes were infected at a multiplicity of infection (MOI) of 10. The cells were transfected with dsRNA 48 hours after infection and harvested for further analysis at 12 after transfection.

### shRNAs, sgRNAs, siRNAs, dsRNA synthesis and transfection

shRNAs targeting mouse *Pntpt1* was synthesized by Tsingke (Kunming, China) and cloned into pLKO.1 vector for lentivirus preparation. The targeted sequence were listed as follows. *shPnpt1-1#*: 5’-CGGCAAAGTTACTGGTGTGAA-3’ and *shPnpt1-2#*: 5’-GGTGGAAGTTTGGCATTAATG-3’.

CRISPR sgRNA sequences used in this study were list as follows. *Dhx9*: 5’-TATGAATCGCTGTGGAGACT-3’ and 5’-GGGTTACCAGCACCAATGGG-3’; *Stat1*: 5’-GTACGATGACAGTTTCCCCA-3’; *p53*: 5’-TCTACAGATGACTGCCATGG-3’; *Mavs*: 5’-TACTAGGTCCCTACCCCTGG-3’ and 5’-GAACAGACCCCAGTGTCATA-3’.

Synthesized siRNAs by RIBOBIO (Guangzhou, China) were used to transiently knockdown the expression of target genes. To examine the effect of target genes on dsRNA induced ISG responses in mouse ESCs, dsRNA was transfected at 48 hours after siRNAs transfection, and 12 hours later, cells were harvested for RT-qPCR analysis. Sequences of siRNAs used in this study were listed as follows: *siDhx9*: 5’-GCTGCCAGAGACTTTGTTA-3’, *siEftud2*: 5’-CCTGTTTCGTAGATTGTTT-3’, *siDdx3x*: 5’-CCTGAACTCTTCAGATAAT-3’, *siDicer1*: 5’-GCGCAAATACAAGCCCTAT-3’, *siTardbp*: 5’-TCAACTATCCCAAAGATAA-3’, *siSf3b3*: 5’-CAGCCATCAAAGAATATGT-3’, *siAdar*: 5’-GCTTCAGCAGATAGAGTTT-3’, *siPrkra*: 5’-GAATCCAATTGGCTCATTA-3’, *siStau1*: 5’-GCAAGAAGATCTCCAAGAA-3’, *siTarbp2*: 5’-ATCTGGATATTGAGGAACT-3’, *siIgf2bp1*: 5’-GCAGGAGATCATGAAGAAA-3’, *siRaly*: 5’-GACAGACCCTGGACATCAA-3’, *siSnrnp200*: 5’-GTGTATGCTCCGAGAGATA-3’, *siRbm39*: 5’-GCACAACAAGCATTACAAA-3’, *siSfpq*: 5’-GGATTTAAAGCCAACTTGT-3’, *siStrbp*: 5’-GATGATAAAGATCCCAACA-3’, *siPrpf19*: 5’-GACACCAACAAGATTCTCA-3’, *siPrpf8*: 5’-GAACATGTCAGGAAGATCA-3’, *siSrsf9*: 5’-GAAGAAACGGTTACGATTA-3’, and *siSrsf10*: 5’-CCAAATCAAATGAAAGCCA-3’.

1k bp *Cas9* dsRNA was generated through *in vitro* transcription as our previously described (*17*). 100 ng dsRNA *per* well (24-well cell plate) was transfected using Lipofectamine 3000 when the cell density was about 80%. 12 hours after dsRNA transfection, cells were harvested for qRT-PCR or Western blot analyses.

### co-immunoprecipitation (co-IP) and Western blotting (WB) assays

Protein-protein or RNA-protein interactions were analyzed by co-immunoprecipitation (co-IP) assays. Briefly, cells were lysed in NP-40 lysis buffer (P0013F, Beyotime) supplemented with complete protease inhibitor cocktail (11697498001, Roche) and RNase inhibitor (N8080119, ThermoFisher Scientific). After centrifugation, the protein concentration of supernatant was determined using a BCA Protein Assay Kit (A55865, Pierce) according to the manufacturer’s instructions. Equal amounts of total protein from the supernatant were pre-cleared with Protein A/G Plus Agarose beads (sc-2003, Santa Cruz Biotechnology) for 1 hour at 4°C. The pre-cleared lysates were then incubated overnight at 4°C with 2 µg of specific primary antibody or an equivalent amount of normal IgG (as a negative control). Protein A/G Plus Agarose beads were added and incubated for an additional 4 hours at 4°C. The beads were collected and washed five times with cold lysis buffer. The bound proteins were eluted by boiling in 2× Laemmli SDS sample buffer (P0015, Beyotime) at 95°C for 10 minutes. The eluted proteins were then resolved by SDS-PAGE and analyzed by immunoblotting with the corresponding antibodies to detect the protein of interest and its putative interaction partners.

For Western blotting (WB) analysis, cells were harvested in RIPA lysis buffer (P0013B, Beyotime) supplemented with protease inhibitors cocktail (11697498001, Roche). Equal amounts of protein were separated by sodium dodecyl sulfate-polyacrylamide gel electrophoresis (SDS-PAGE) and then electrophoretically transferred onto polyvinylidene difluoride (PVDF) membranes (88520, ThermoFisher Scientific). After transfer, the membranes were blocked with 5% bovine serum albumin (BSA) (A1933, Sigma-Aldrich) in Tris-buffered saline containing 0.1% Tween-20 (TBST) for 1 hour at room temperature. Subsequently, the membranes were incubated overnight at 4°C with specific primary antibodies diluted in the blocking solution. Following three washes (10 min each) with TBST, the membranes were incubated with the corresponding horseradish peroxidase (HRP)-conjugated secondary antibodies for 1 hour at room temperature. After another series of TBST washes, protein bands were visualized using an enhanced chemiluminescence (ECL) detection kit (34577, ThermoFisher Scientific) and imaged with a ChemiDoc™ MP Imaging System (Bio-Rad).

The following antibodies were used for WB or IP analysis: anti-Tubulin (1:5000 for WB, 66031-1-Ig, Proteintech), anti-Actin (1:5000 for WB, 66009-1-Ig, Proteintech), anti-p53 (1:5000 for WB and 2 µg for each IP, 10442-1-AP, Proteintech), anti-p53 (1:1000 for WB, ab77813, Abcam), anti-Puromycin (1:1000 for WB, MAB3343, Sigma-Aldrich), anti-Stat1 (1:1000 for WB and 2 µg for each IP, 14994S, Cell Signaling Technology), anti-p701-Stat1 (1:1000 for WB, 9167S, Cell Signaling Technology), anti-Dhx9 (1:5000 for WB and 2 µg for each IP, 17721-1-AP, Proteintech), anti-Mdm2 (1:1000 for WB, 66511-1-Ig, Proteintech), anti-Ubiquitin (1:1000 for WB, sc-8017, Santa Cruz Biotechnology), anti-Cul4A (1:1000 for WB, 14851-1-AP, Proteintech), anti-Ddb1 (1:1000 for WB, PA5-21282, Invitrogen), anti-Ddb2 (1:1000 for WB, ab77765, Abcam), anti-Rbx1 (1:1000 for WB, 14895-1-AP, Proteintech), anti-Gapdh (1:5000 for WB, 60004-1-Ig, Proteintech), anti-J2 (2 µg for each IP, 76651L, Cell Signaling Technology), anti-Tcf4 (1:1000 for WB, 05-511, Millipore), mouse control IgG (2 µg for each IP, B900620, Proteintech), and rabbit control IgG (2 µg for each IP, B900610, Proteintech); HRP conjugated anti-mouse (1:5000 for WB, 31430, Invitrogen) or rabbit (1:5000 for WB, 31460, Invitrogen) IgG (H+L) was used as the secondary antibody for WB analysis.

### Cell immunofluorescence (IF) assays

Mouse or human ESC cells were seeded in 24-well plate and transfected with dsRNA or not as indicated. Twenty-four hours after transfection, cells were fixed with 4% paraformaldehyde and then permeabilized with 0.2% Triton-X100. After blocking with 2% BSA (A1933, Sigma-Aldrich), cells were incubated with the indicated primary antibodies. Alexa Fluor 555 or 488 dye conjugated secondary antibodies were used to detect the primary antibody. Nuclei were stained with 4,6-diamidino-2-phenylindole (DAPI) (1:5000, D9542, Sigma-Aldrich) was used to visualize DNA in nucleus. The following primary antibodies were used in this study for IF staining: anti-J2 (1:300, 10010500, Scicons), anti-Dhx9 (1:300, ab26271, Abcam), anti-Stat1 (1:100, ab239360, Abcam), anti-p53 (1:100, ab246550, Abcam), anti-Ddb1 (1:100, PA5-21282, ThermoFisher Scientific), anti-Mdm2 (1:100, 66500-1-Ig, Proteintech), anti-Cul4A (1:100, 14851-1-AP, Proteintech). Micrographs were taken with the LSM880 confocal microscope (Car Zeiss, Germany).

### Total RNA extraction, and qRT-PCR

Total RNA from cells or zebrafish embryos were extracted using TRIzol reagents (DP424, TianGen) according to the manufacturer’s instructions. Reverse transcription was carried out by using a First-Strand cDNA synthesis kit (K1622, ThermoFisher Scientific). Expression of target genes were quantified using the LightCycler® 480 SYBR Green I Master kit (04707516001, Roche) on a LightCycler480 system (Roche). All reactions were run in replicates of at least of four. The expression of mouse *Actin* or human *ACTIN* was used as a loading control respectively. Primer used were listed as follows: mouse *β-Actin*: forward, 5’- TGAGCGCAAGTACTCTGTGTGGAT-3’ and reverse, 5’-ACTCATCGTACTCCTGCTTGCTGA-3’; mouse *Isg15*: forward, 5’-GGTGTCCGTGACTAACTCCAT-3’ and reverse, 5’- TGGAAAGGGTAAGACCGTCCT-3’; mouse *Cxcl10*: forward, 5’- CCAAGTGCTGCCGTCATTTTC -3’ and reverse, 5’- GGCTCGCAGGGATGATTTCAA -3’; mouse *Oasl1*: forward, 5’-CAGGAGCTGTACGGCTTCC-3’ and reverse, 5’-CCTACCTTGAGTACCTTGAGCAC-3’; mouse *Ifih1*: forward, 5’- AGATCAACACCTGTGGTAACACC-3’ and reverse, 5’-CTCTAGGGCCTCCACGAACA-3’; mouse *Pmaip1*: forward, 5’-GCAGAGCTACCACCTGAGTTC-3’ and reverse, 5’- CTTTTGCGACTTCCCAGGCA-3’; mouse *Tp53inp1*: forward, 5’- AAGTGGTCCCAGAATGGAAGC-3’ and reverse, 5’-GGCGAAAACTCTTGGGTTGT-3’; human *β-ACTIN*: forward, 5’-ACCGAGCGCGGCTACAG-3’ and reverse, 5’- CTTAATGTCACGCACGATTTCC-3’; human *ISG15*: forward, 5’-CGCAGATCACCCAGAAGATCG-3’ and reverse, 5’-TTCGTCGCATTTGTCCACCA-3’; human *CXCL10*: forward, 5’-GTGGCATTCAAGGAGTACCTC-3’ and reverse, 5’-TGATGGCCTTCGATTCTGGATT-3’; human *IFIH1*: forward, 5’-TCGAATGGGTATTCCACAGACG-3’ and reverse, 5’-GTGGCGACTGTCCTCTGAA-3’; human *OASL1*: forward, 5’-CTGATGCAGGAACTGTATAGCAC-3’ and reverse, 5’-CACAGCGTCTAGCACCTCTT-3’; human *PMAIP1*: forward, 5’-ACCAAGCCGGATTTGCGATT-3’ and reverse, 5’-ACTTGCACTTGTTCCTCGTGG-3’; human *TP53INP1*: forward, 5’-TTCCTCCAACCAAGAACCAGA-3’ and reverse, 5’-GCTCAGTAGGTGACTCTTCACT-3’; mouse *Pnpt1*: forward, 5’-CCTTCTCCATATTTGCACCAACA-3’ and reverse, 5’-TGTCGCGGTATAAACTGCTCC-3’; mouse *Dhx9*: forward, 5’-CCGAGGAGCCAACCTTAAAGA-3’ and reverse, 5’-TGTCCAATTTCCATGAAGCCC-3’; mouse *Eftud2*: forward, 5’-GATCGAGCATACCTACACTGGC-3’ and reverse, 5’-GTACATCTTCGTCGTGTGGCA-3’; mouse *Igf2bp1*: forward, 5’-CGGCAACCTCAACGAGAGT-3’ and reverse, 5’-GTAGCCGGATTTGACCAAGAA-3’; mouse *Tarbp2*: forward, 5’- GGCTCCGGCACTACTACAG-3’ and reverse, 5’-TGGTCCCATACTCCTGAAGAAG-3’; mouse *Ddx3x*: forward, 5’-CAGAGTGGAGGAAGTACAGCA-3’ and reverse, 5’-TCACCCCGTGATCCAAAACTG-3’; mouse *Raly*: forward, 5’-ATCGGCAATTTGAATACAGCCA-3’ and reverse, 5’-GACAGGGGTCTCTTACTTCCA-3’; mouse *Snrnp200*: forward, 5’-AGTTTGCAGTACGAGTACAAGG-3’, and reverse, 5’-CCCTCCAACTTCCCAACCAG-3’; mouse *Dicer1*: forward, 5’-GGTCCTTTCTTTGGACTGCCA-3’, and reverse, 5’-GCGATGAACGTCTTCCCTGA-3’; mouse *Rbm39*, forward, 5’- CATGCTTGAGGCCCCTTACAA-3’, and reverse, 5’-TTCCGCTCTCGACTTTTGCTC-3’; mouse *Sfpq*, forward, 5’-TGTCGGTTGTTTGTGGGGAAT-3’, and reverse, 5’-AACCCGAACCCTTTGCCTTT-3’; mouse *Strbp*, forward, 5’-GTGTGGTGTAATGAGGATTGGC-3’, and reverse, 5’-TTTCCGTGGGTTTGTCTTTGC-3’; mouse *Tardbp*: forward, 5’-AATCAGGGTGGGTTTGGTAACA-3’, and reverse, 5’-GCTGGGTTAATGCTAAAAGCAC-3’; mouse *Sf3b3*: forward, 5’-TCCTTCAGGACAGTTGAATGAGT-3’, and reverse, 5’-GTCTGATGGGTCGAGGGAGAT-3’; mouse *Prpf19*: forward, 5’-CCTCACAGTGGCCTACCCT-3’, and reverse, 5’-CAGTTGCTGGCGAAGAGTGAA-3’; mouse *Adar*: forward, 5’-TGAGCATAGCAAGTGGAGATACC-3’, and reverse, 5’-GCCGCCCTTTGAGAAACTCT-3’; mouse *Prpf8*: forward, 5’-CCGCTGCCGGATTATATGTCA-3’, and reverse, 5’-TCCTTCTGAGCATCCACAAAC-3’; mouse *Srsf9*: forward, 5’-CAACCTTCCGTCCGACGTG-3’, and reverse, 5’-GGCCATAATCGTAACCGTTTCTT-3’; mouse *Srsf10*: forward, 5’-CGAAGCCGGAGTTATGAAAGG-3’, and reverse, 5’-GTCGGTCTACTGTTTCTAGGACT-3’; mouse *Prkra*: forward, 5’-CAGCGGGACCTTCAGTTTG-3’, and reverse, 5’-GCACATCGGATCTTTCACATTCA-3’; mouse Stau1: forward, 5’-GGACCCTCACTCTCGGATG-3’, and reverse, 5’-TTCTGGCAGGGGTTCACTCT-3’; *Zika virus MR766*: forward, 5’-TTGGTCATGATACTGCTGATTGC-3’, and reverse, 5’-CCCTCCACGAAGTCTCTATTGC-3’; zebrafish *gapdh*: forward, 5’-TTGCCGTTCATCCATCTTTG-3’, and reverse, 5’-TGCTGTAACCGAACTCATTGTC-3’; zebrafish *phlda3*: forward, 5’-TTCGAGCAGATGACCACTCT-3’ and reverse, 5’-CTCAAAGTTCGACGTCACCT-3’; zebrafish *cdkn1a*: forward, 5’-ATGGCGGCGCACAAGCGGAT-3’ and reverse, 5’-CACTAGACGCTTCTTGGCTT-3’; zebrafish *isg15*: forward, 5’-TGACGATGCAGCTGACTGTA-3’ and reverse, 5’-CAACTTCATGCCAGACTCGA-3’; zebrafish *casp8*: forward, 5’-TTCTGCGACTGGATGAGCAA-3’ and reverse, 5’-TCCATATCAGTGCCTGTTCG-3’; zebrafish *ifit8*: forward, 5’-AGAGGAGTTCACGCTGAAGAA-3’ and reverse, 5’-TGTGTTCCACTCCTTGTCAT-3’; zebrafish *cxcl12b*: forward, 5’-GCATGGATAGCAAAGTAGTAG-3’ and reverse, 5’-AGTGTGAGACTCCAGGACAC-3’; zebrafish *ifng*: forward, 5’-GACATACAGTGAAGCCAGTG-3’ and reverse, 5’-CTCTATAGACACGCTTCAGC-3’; zebrafish *ifnphi1*: forward, 5’-CTCTGCGTCTACTTGCGAAT-3’ and reverse, 5’- TTCACAACTCTCCGGATCTG-3’; zebrafish *ifnphi2*: forward, 5’-TGCGTTCTTATGTCCAGCAC-3’ and reverse, 5’-TTCCTTGAGCTCTCATCCTC-3’; zebrafish *ifnphi3*: forward, 5’-GAGGACCTATACACTTCTGG-3’ and reverse, 5’-AGAGTCGAGGATATTGTCTC-3’; zebrafish *ifnphi4*: forward, 5’-TCCATGATGACAATGAGGAC-3’ and reverse, 5’-AGGAGTGCAGCTGATTCATC-3’.

### RNA-sequencing analysis

Total RNA was extracted from 6 hpf zebrafish embryos or the indicated cells as described above. Poly(A)^+^ mRNA was isolated using the Poly(A) mRNA Magnetic Isolation Module (NEB). Libraries were prepared using the NEBNext Ultra RNA Library Prep Kit for Illumina according to the manufacturer’s instructions. Briefly, mRNA was fragmented and reverse-transcribed to generate first- and second-strand cDNA, followed by purification with AxyPrep Mag PCR Clean-up beads (Axygen). cDNA ends were repaired, A-tailed, and ligated to sequencing adapters. Fragments of ∼300 bp were size-selected and amplified by 11 cycles of PCR. Paired-end sequencing (150 bp) was performed on an Illumina HiSeq 2000 platform.

Raw reads were processed using Cutadapt and FastQC. Reads from three biological replicates were assembled and aligned using StringTie. Differential gene expression was analyzed using DESeq2 and edgeR. Volcano plots were generated with GraphPad Prism 9, and heatmaps were produced using TBtools (*58*). Genes shown in the heatmap of Fig. 7B were selected based on the following criteria: expression >0.3 FPKM and ≥21.8-fold induction in wild-type embryos after dsRNA injection, with no significant change observed in MZ*tp53* mutants. Top 100 upregulated genes by dsRNA in wild-type mESCs were selected and shown in the heatmap of Fig 1C. Top 100 upregulated genes and the upregulated TP53 target genes by dsRNA in wild-type hESCs were selected and shown in the heatmap of Fig. 6B. ISG and P53 target genes were selected and highlighted as previously reported (*59*, *60*).

### Cleavage Under Targets and Tagmentation (CUT&Tag) assays

CUT&Tag assays were performed as previously described using the NovoNGS CUT&Tag 4.0 High-Sensitivity Kit (N259-YH01, Novoprotein) according to the manufacturer’s instructions with minor modifications (*61*). Briefly, 12 hours after dsRNA transfection, 1×10^6^ mouse ESC cells were harvested and bound to ConA beads. Cells were permeabilized with digitonin and incubated with p53 (10442-1-AP, Proteintech), Stat1 (14994S, Cell Signaling Technology), or Dhx9 (17721-1-AP, Proteintech) primary antibody overnight at 4°C. After washing, cells were incubated with the secondary antibody (B900610, Proteintech) for 1 hour at room temperature, followed by incubation with pAG-Tn5 transposase for 1 hour. Tagmentation was performed at 37°C for 1 hour and stopped by adding proteinase K (10401ES80, Yeasen). DNA was purified using KAPA Pure Beads (07983280001, Roche) and amplified by PCR. Libraries were sequenced on an illumina NovaSeq 6000 platform with 150 bp paired-end reads.

Sequencing reads were aligned to reference genome (GRCm38/mm10) using Bowtie2 (v2.4.2) with parameters “--end-to-end --very-sensitive --no-mixed --no-discordant”. Duplications were removed using Picard MarkDuplicates. Peaks were called using MACS2 (v2.2.7.1) with a *q*-value cutoff of 0.05 and without an input control. BigWig files for visualization were generated using deepTools (v3.5.1) bamCoverage with RPKM normalization. Genome browser tracks were visualized with the Integrative Genomics Viewer (IGV).

### *In vivo* ubiquitination assays

To examine the ubiquitination of endogenous p53 or Stat1 in mouse ESC cells, cells were harvested at 12 hours after dsRNA transfection and *in vivo* ubiquitination assays were carried out as previously described (*62*). Briefly, cells were lysed in an SDS lysis buffer (50 mM Tris-HCl [pH 6.8], 1.5% SDS) at 95°C for 15 min. Following 10-fold dilution of the lysate with extraction buffer C (EBC)-bovine serum albumin (BSA) (50 mM Tris-HCl [pH 6.8], 180 mM NaCl, 0.5% NP-40, and 0.5% BSA) plus protease inhibitors (11697498001, Roche), the cell lysates were immunoprecipitated with anti-p53 (10442-1-AP, Proteintech) or Stat1 (14994S, Cell Signaling Technology) antibody. The bound proteins were eluted with 2× SDS loading buffer at 95°C for 5 min. Western blot analysis was performed using an anti-Ubiquitin (sc-8017, Santa Cruz Biotechnology) antibody.

### dsRNA pull down and mass spectrometry (MS)

24 hours after dsRNA transfection, mouse ESC cells were harvested and lysazed for 30 min on ice in the lysis buffer (50 mM Tris [pH 7.5], 150 mM NaCl, 1 mM ethylenediaminetetraacetic acid [EDTA], and 0.2% Nonidet P-40, 1.5 mM MgCl_2_) supplemented with RNase inhibitor (N8080119, ThermoFisher Scientific), and protease inhibitor cocktail (11697498001, Roche). Cell lysates were clarified by centrifugation at 16,000 × *g* for 15 min and the protein concentrations were determined with the BCA assay. Next, equal amount of extracted cellular protein was incubated with dynabeads (10004D, ThermoFisher Scientific) coupled with control IgG or J2 antibody overnight with gently rotation at 4°C. Finally, the beads were washed with the lysis buffer for five times and then eluted with 2 × SDS loading buffer at 95°C for 15 min. The purified proteins were quantitatively analyzed by liquid chromatography-tandem mass spectrometry or analyzed by WB assays. For the liquid chromatography-tandem mass spectrometry data analysis, proteins present in at least two replicates were retained and statistical cut-offs (*p* value < 0.01 and fold change > 2) were applied to define the potential dsRNA binding proteins.

### Quantification and statistical analysis

All experiments were repeated at least three times with similar results. Data plotting and statistical tests were performed using GraphPad Prism 9. Statistical information is described in the figure legends.

## Acknowledgments

We thank professors Yuan Hu (Chongqing medical university) for the plasmid encoding human Dhx9, and Bo Zhao (Kunming Institute of Zoology, Chinese Academy of Sciences) for the human ESC line. We thank the staff members of the National Research Facility for Phenotypic & Genetic Analysis of Model Animals (Primate Facility) (https://cstr.cn/31137.02.NPRC) and the Institutional Center for Shared Technologies and Facilities of KIZ for technical support.

## Author contributions

B.M. and M.S. conceptualized the project; P.M., T.L. and J.X. designed the methodology; P.M., J.X., Y.L., R.L., and T.L. performed the investigation; Y.Z., X.Y., and J.L. provided facility support and reagents; P.M., J.X., and T.L. visualized the datasets; P.M., B.M., and M.S. acquired the funding; B.M. administrated the project; B.M. supervised the project; P.M., J.X. and B.M. wrote the original draft; P.M., J.X., B.M., and M.S. reviewed and edited the manuscript.

## Data, code, and materials availability

Further information and requests for resources and reagents should be directed to and will be fulfilled by the corresponding author. All biological materials are available upon request from the corresponding author and signing of appropriate materials transfer agreements.

## Supplementary Materials

**Figure S1.**
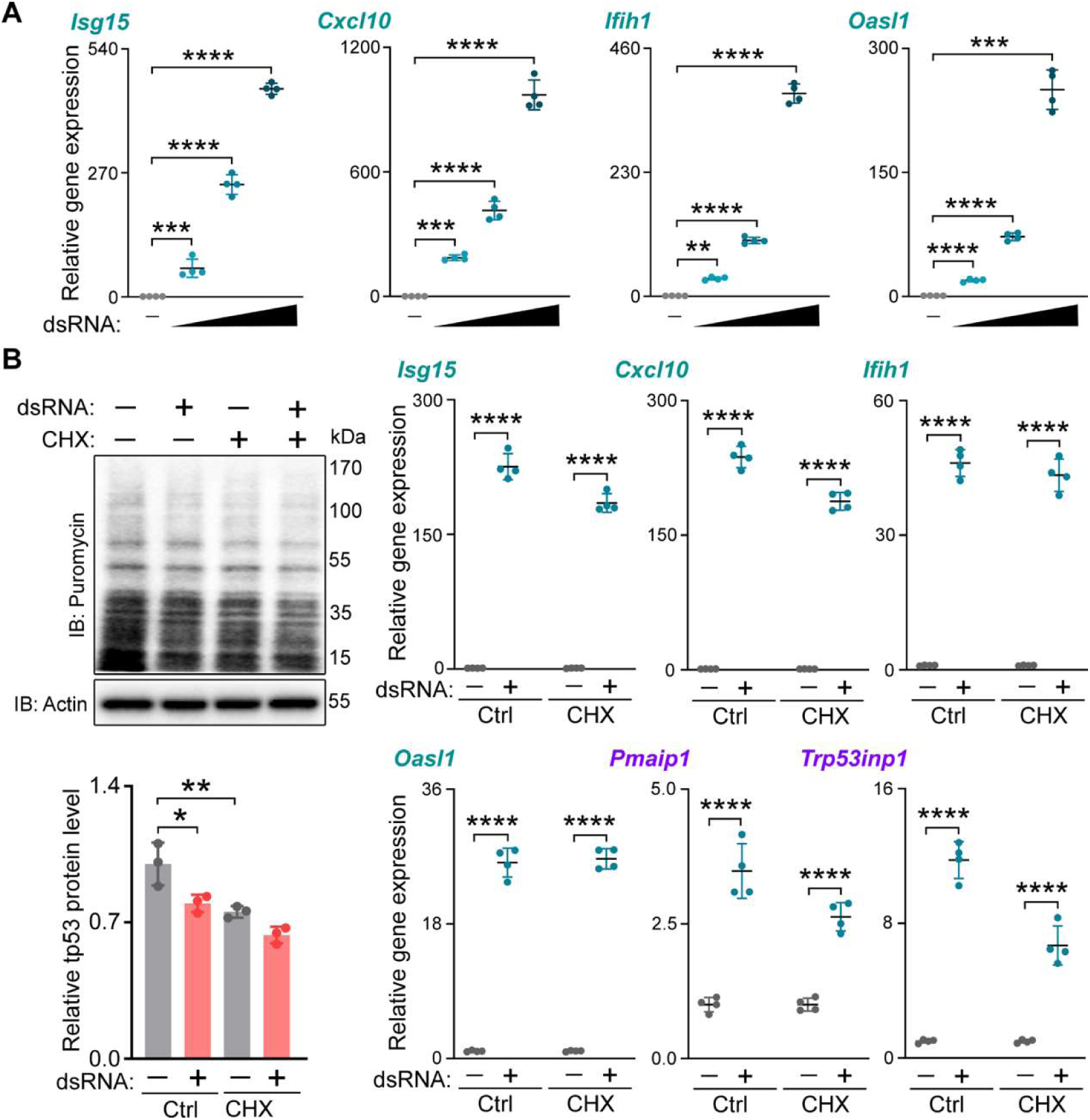
dsRNA induced an ISG response in mESCs which is not dependent on protein synthesis. **(A)** RT-qPCR results showing relative expressions of the indicated ISGs (*Isg15*, *Cxcl10*, *Ifih1*, and *Oasl1*) in mESCs with 0, 50, 100, and 200 ng/well (24-well plate) dsRNA transfection. Ordinary one-way ANOVA with multiple comparisons. **, *p* < 0.01; ***, *p* < 0.001; ****, *p* < 0.0001. **(B)** Western blotting results showing the translational inhibition by CHX treatment and RT-qPCR results showing relative expressions of the indicated ISGs (*Isg15*, *Cxcl10*, *Ifih1*, and *Oasl1*) and p53 target genes (*Pmaip1* and *Trp53inp1*) in control and dsRNA transfected mESCs with CHX treatment or not. Ctrl, control; CHX, cycloheximide; IB, immunoblotting; Ordinary one-way ANOVA with multiple comparisons. *, *p* < 0.05; **, *p* < 0.01; ****, *p* < 0.0001.

**Figure S2.**
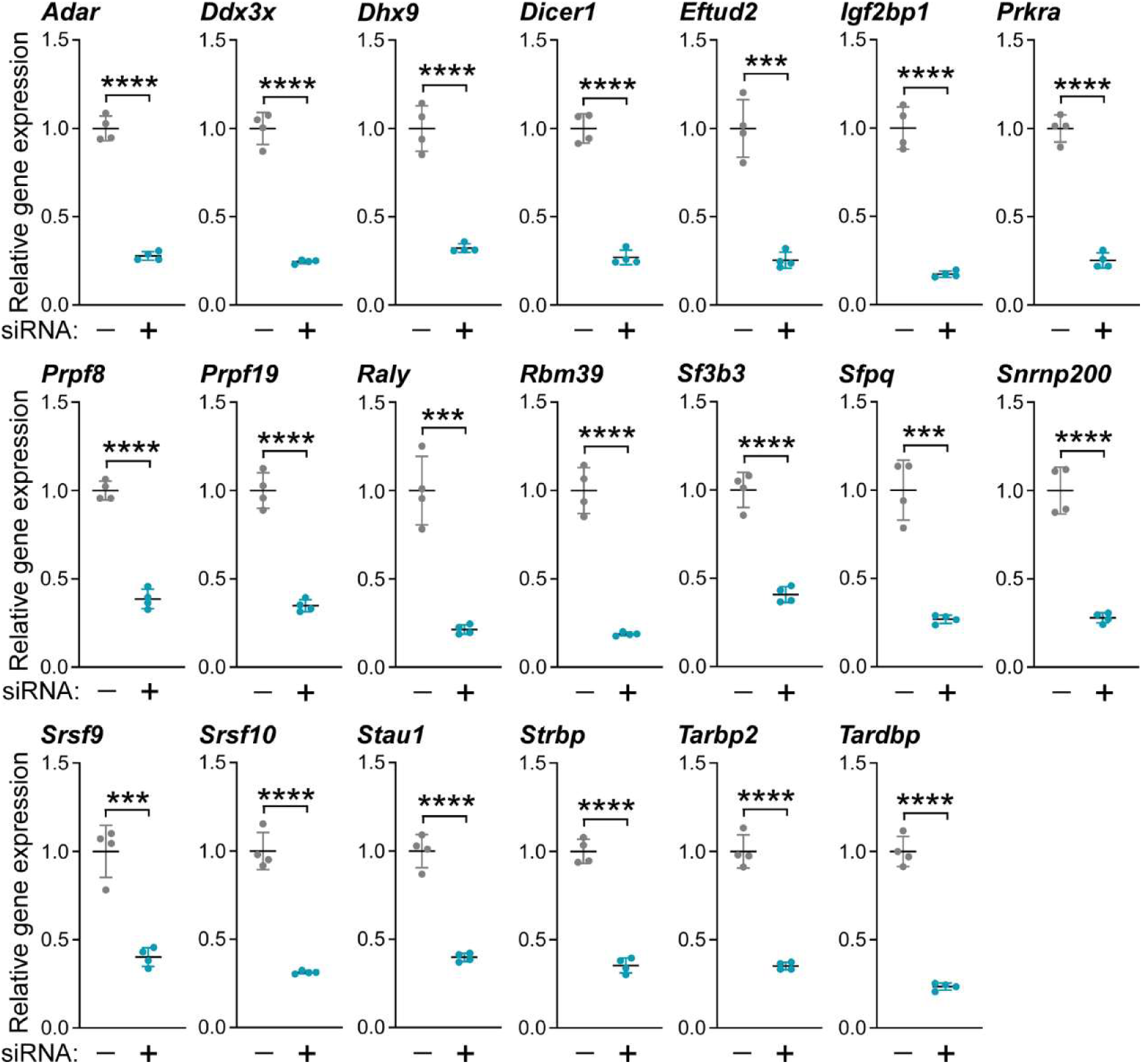
siRNA-mediated knockdown of the indicated RNA binding proteins in mESCs. RT-qPCR results showing relative expressions of the indicated genes in mESCs transfected with their siRNAs. Student’s *t*-test. ***, *p* < 0.001; ****, *p* < 0.0001.

**Figure S3.**
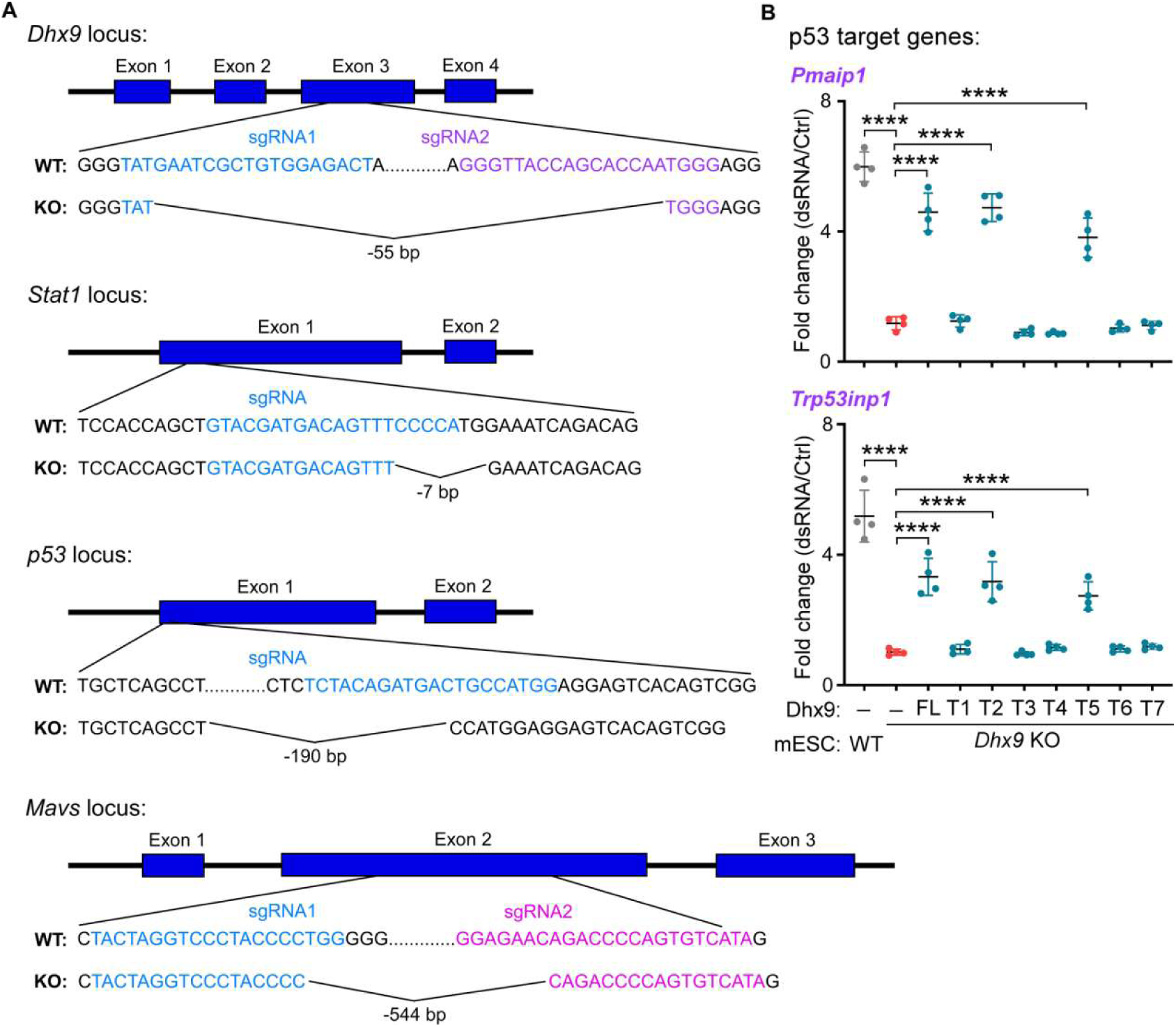
Strategy for *Dhx9*, *Stat1*, *p53*, and *Mavs* knockout in mESCs and the expression of p53 target genes in *Dhx9* KO mESCs with the indicated Dhx9 truncates overexpression. **(A)** Schematic representation of gRNA mediated gene knockout of *Dhx9*, *Stat1*, *p53, and Mavs* genes in mESCs. WT, wild type; KO, knockout. **(B)** RT-qPCR results showing expressional fold changes of the indicated p53 target genes (*Pmaip1* and *Trp53inp1*) in dsRNA transfected *vs* control mESCs of WT or *Dhx9* KO with overexpression of the indicated Dhx9 truncates shown in Fig. 2C. Ordinary one-way ANOVA with multiple comparisons. ****, *p* < 0.0001. WT, wild type; KO, knockout; T, truncate; Ctrl, control.

**Figure S4.**
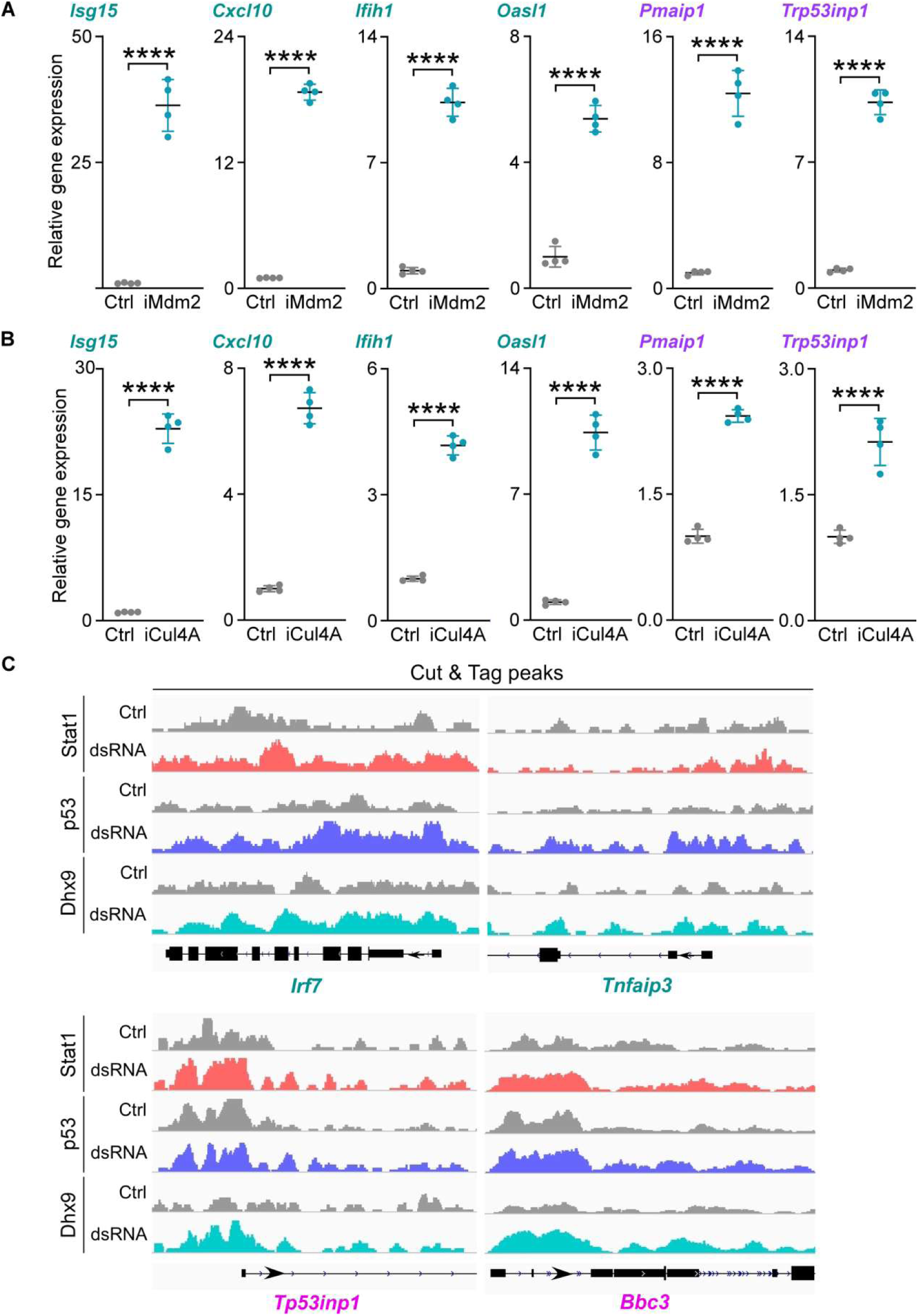
Mdm2 or Cul4A inhibition induces an ISG response and p53 target gene activation in mESCs, and Dhx9, p53, and Stat1 cooperate to activate ISGs and p53 target genes. **(A, B)** RT-qPCR results showing relative expression levels of *ISG15* in Nutlin (Mdm2 inhibitor, iMdm2) (A) or KH-4-43 (Cul4A inhibitor, iCul4A) (B) treated mESCs. Ctrl, control; Student’s *t*-test. ****, *p* < 0.0001. **(C)** Cut&Tag peaks (visualized by IGV) of Dhx9, p53, and Stat1 for the selected ISG genes (*Irf7* and *Tnfaip3*) or p53 target genes (*Trp53inp1 and Bbc3*) in control or dsRNA transfected WT mESCs. Ctrl, control.

**Figure S5.**
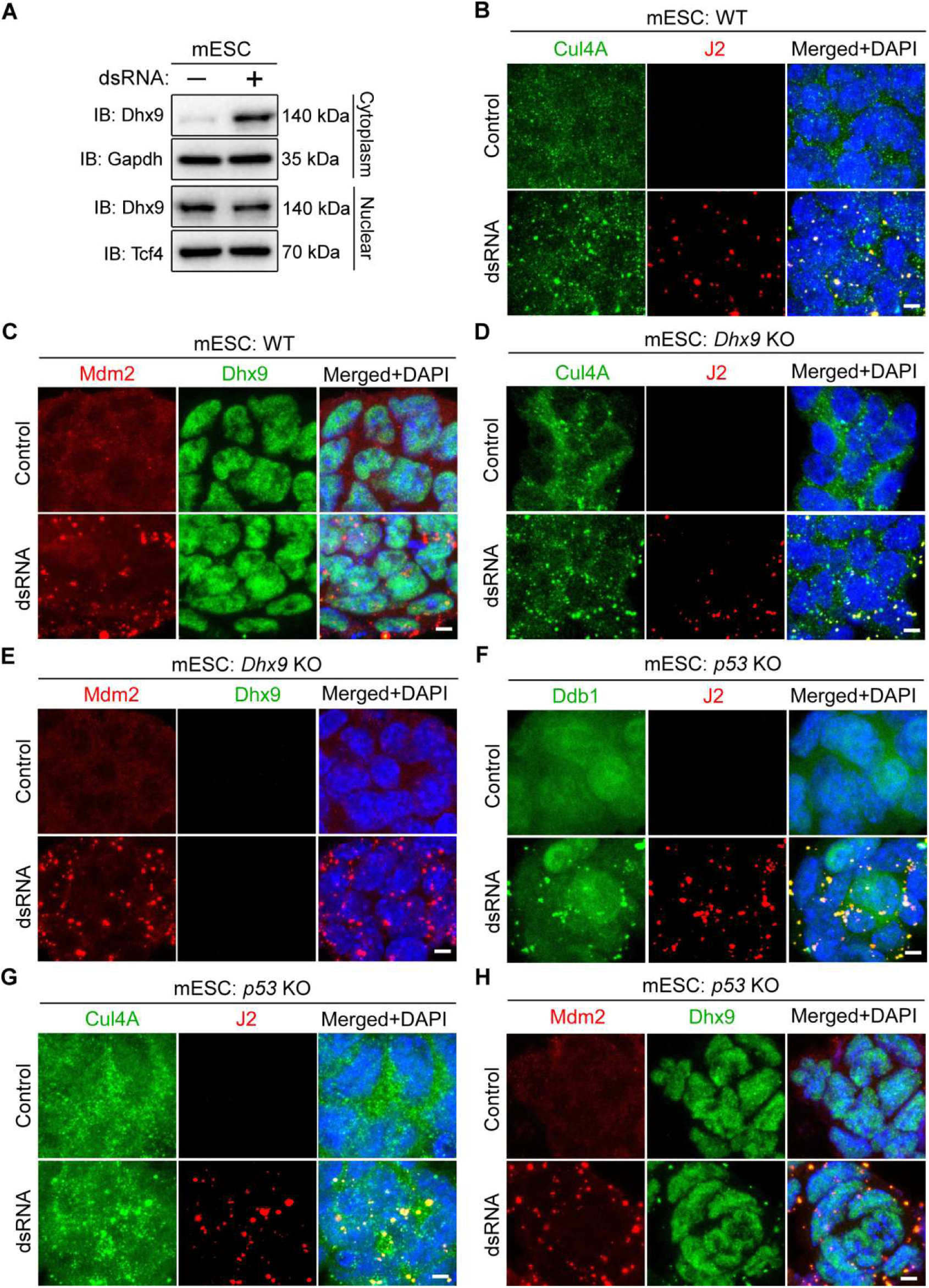
Subcellular localization of Dhx9 in dsRNA transfected mESCs and dsRNA (J2) co-staining with Dhx9, Mdm2, Cul4A, or Ddb1 in WT, *Dhx9* KO, or *p53* KO mESCs. **(A)** Immunoblotting images showing the subcellular expressions of Dhx9 protein in control or dsRNA transfected mESCs. IB, immunoblotting. **(B-H)** Representative immunofluorescence images showing the co-staining of dsRNA (J2) and Cul4A (B, D, and G), Mdm2 (C, E, and H), or Ddb1 (F) in WT, *Dhx9* KO, or *p53* KO mESCs. WT, wild type; KO, knockout. Scale bar, 5 μm.

**Figure S6.**
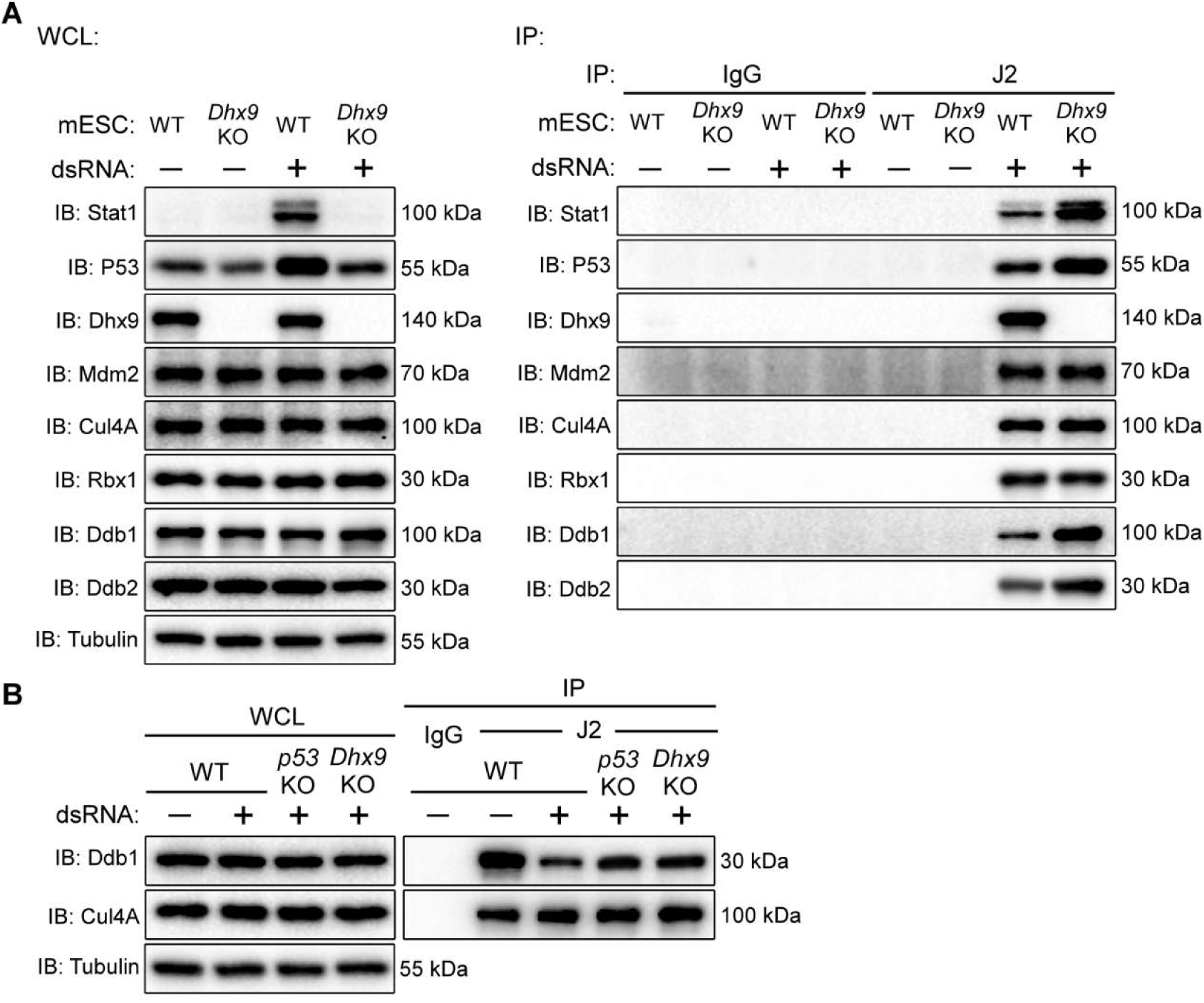
The Dhx9 mediated dsRNA sensory mechanism to stabilizes Stat1 and p53 proteins for ISG response in mESCs. **(A)** dsRNA pull down assays showing the protein components of dsRNA positive foci in WT and *Dhx9* KO mESCs. WCL, whole cell lysate; IP, immunoprecipitation; IB, immunoblotting. **(B)** Co-IP assays showing the interactions between Ddb1 and Cul4A in control or dsRNA transfected mESCs of WT, *Dhx9* KO, and *p53* KO. WCL, whole cell lysate; IP, immunoprecipitation; IB, immunoblotting.

**Figure S7.**
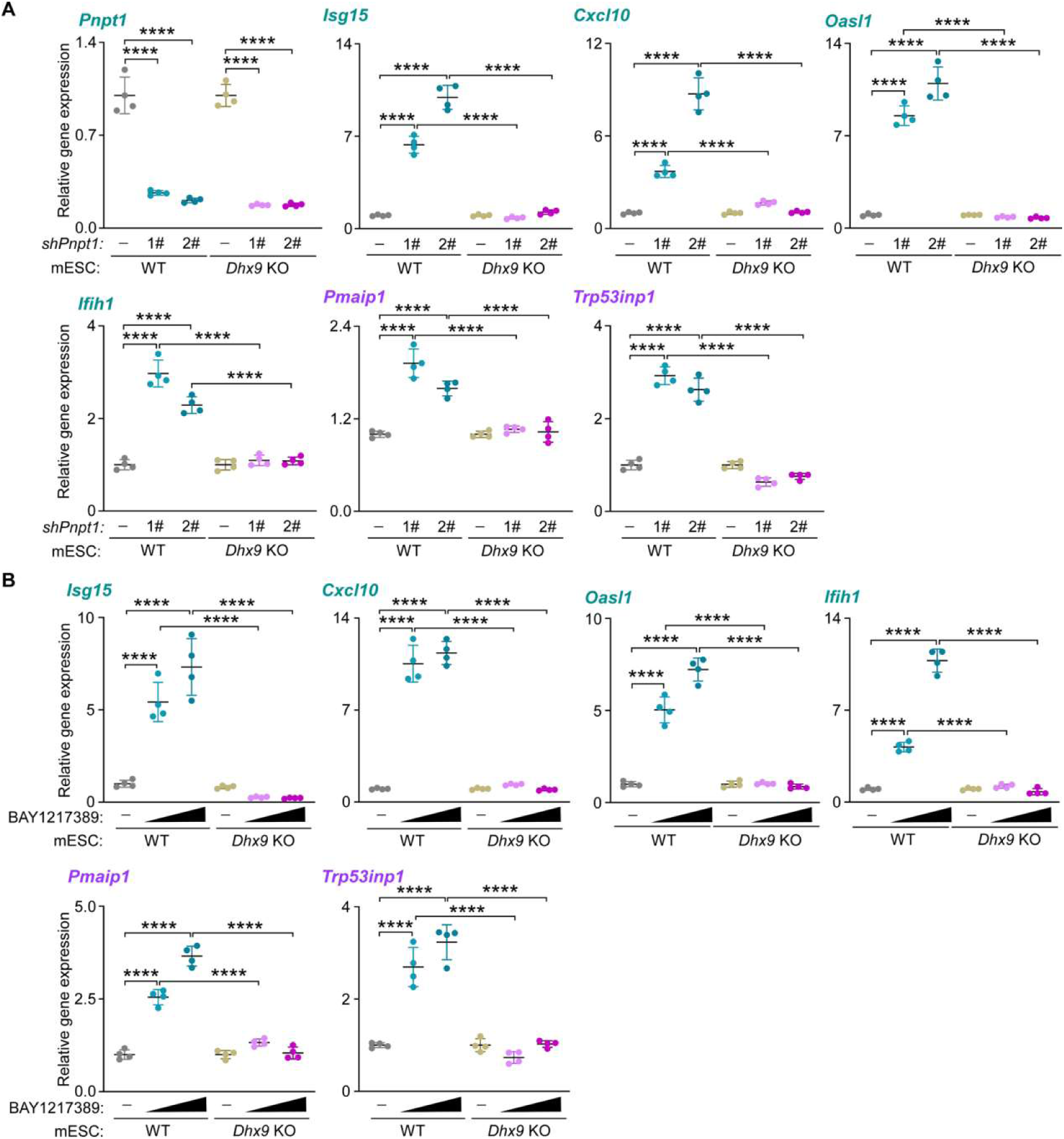
The dsRNA-Dhx9-p53-Stat1 cascade functions in mESCs under dsRNA stress. **(A)** RT-qPCR results showing the relative expression level of *Pnpt1* and the indicated ISGs (*Isg15*, *Cxcl10*, *Oasl1* and *Ifih1*) and p53 target genes (*Pmaip1* and *Trp53inp1*) in WT or *Dhx9* KO mESCs with *Pnpt1* knockdown by two independent shRNAs. Ordinary one-way ANOVA with multiple comparisons. n.s., not significant; ****, *p* < 0.0001. WT, wild type; KO, knockout. **(B)** RT-qPCR results showing the relative expression level of the indicated ISGs (*Isg15*, *Cxcl10*, *Oasl1* and *Ifih1*) and p53 target genes (*Pmaip1* and *Trp53inp1*) in WT or *Dhx9* KO mESCs treated with BAY1217389 at the concentration of 10 and 50 mM. Ordinary one-way ANOVA with multiple comparisons. n.s., not significant; ****, *p* < 0.0001. WT, wild type; KO, knockout.

**Figure S8.**
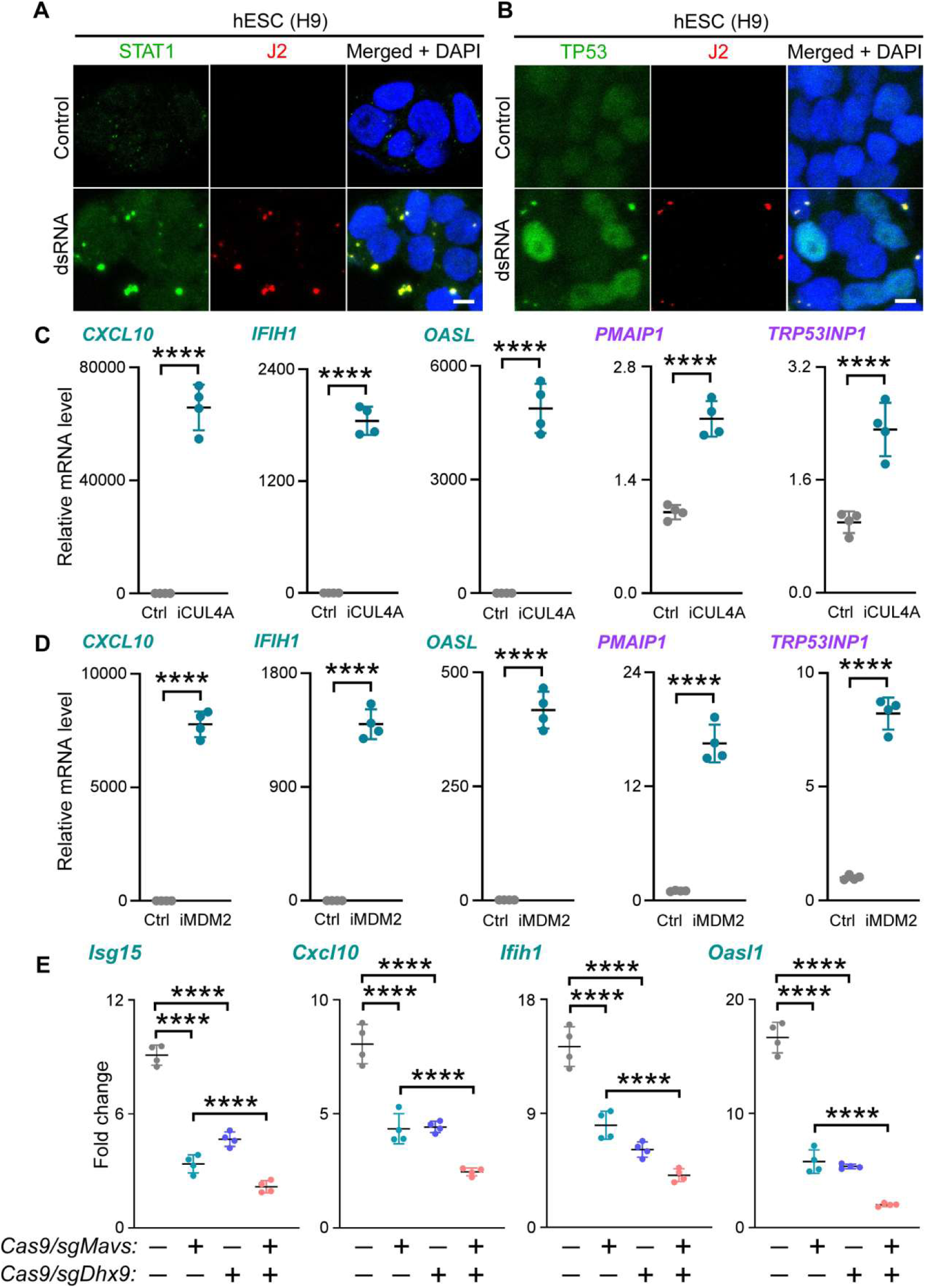
Dhx9-mediated dsRNA signaling in hESCs and differentiated N2a cells. **(A, B)** Representative immunofluorescence images showing the co-staining of dsRNA (J2) and STAT1 (A) and TP53 (B) in hESCs. Scale bar, 5 μm. **(C, D)** RT-qPCR results showing relative expression levels of the indicated ISGs (*CXCL10*, *IFIH1*, and *OASL*) and TP53 target genes (*PMAIP1* and *TRP53INP1*) in Nutlin (MDM2 inhibitor, iMDM2) (C) or KH-4-43 (CUL4A inhibitor, iCUL4A) (D) treated hESCs. Student’s *t*-test. ****, *p* < 0.0001. Ctrl, control. **(E)** RT-qPCR results showing relative expression levels of the indicated ISGs (*Isg15*, *Cxcl10*, *Ifih1*, and *Oasl1*) and p53 target genes (*Pmaip1* and *Trp53inp1*) in N2a cells with sgRNAs against *Mavs* or *Dhx9* transfection. Ordinary one-way ANOVA with multiple comparisons. ****, *p* < 0.0001.

**Figure S9.**
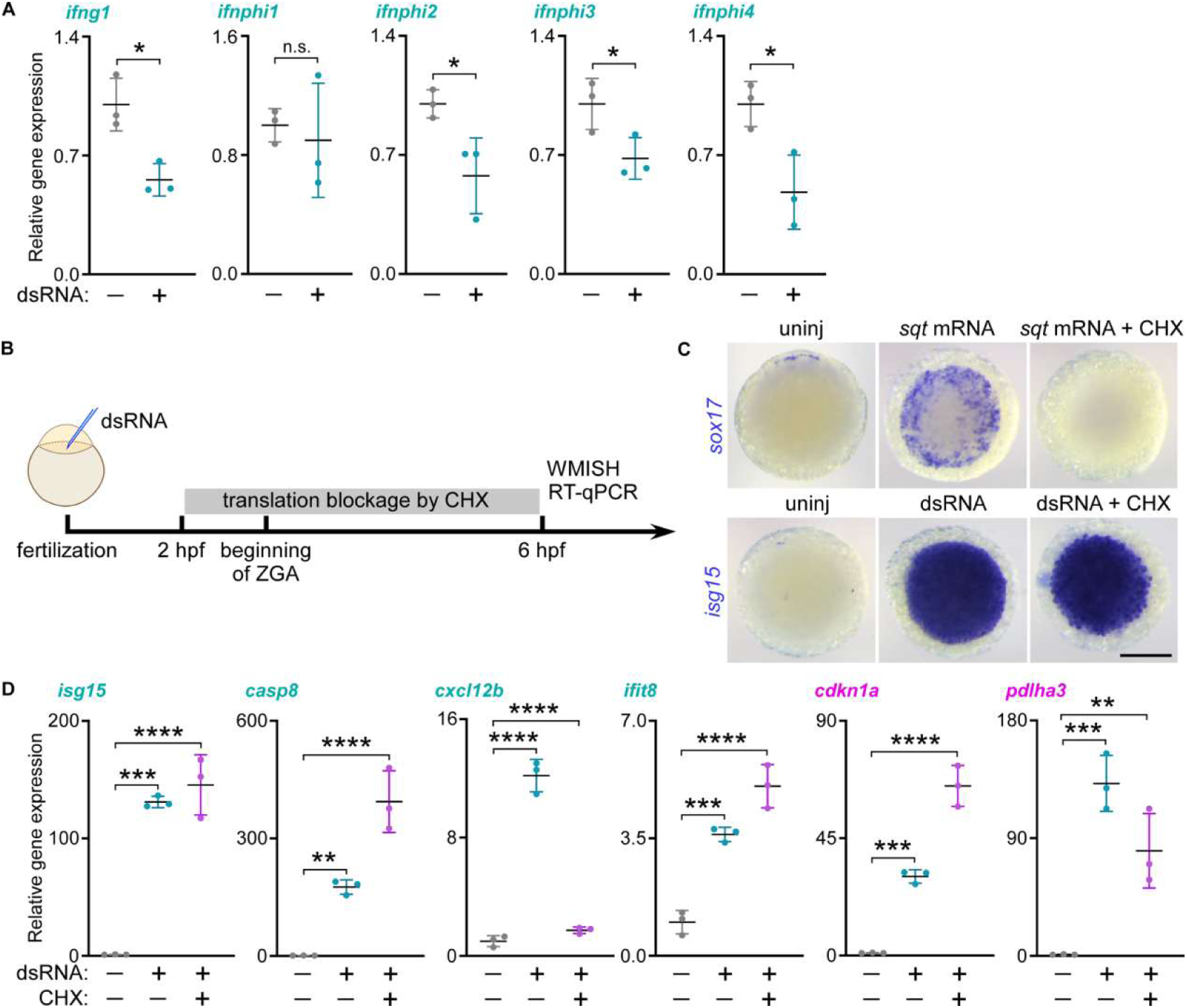
dsRNA induced ISG response and tp53 target genes activation independent on new protein translation in zebrafish early embryos. **(A)** RT-qPCR results showing relative expression levels of the indicated IFN ligands (*ifng1, ifnphi2, ifnphi3, ifnphi4, and ifnphi5*) in control or dsRNA injected zebrafish embryos at 6 hpf. Student’s *t*-test; n.s., not significant; *, *p* < 0.05. **(B-D)** Schematic image showing the experiment strategy (B). WMISH showing the expression of sox17 and isg15 in zebrafish embryos at 6 hpf of control, *sqt* mRNA/dsRNA injection, or *sqt* mRNA/dsRNA injection with CHX treatment (C). Scale bar, 200 μm. RT-qPCR results showing relative expression levels of the indicated ISGs (*isg15, casp8, cxcl12b,* and *ifit8*) and tp53 target genes (*cdkn1a* and *pdlha3*) in control or dsRNA injected zebrafish embryos with CHX treatment or not at 6 hpf. Ordinary one-way ANOVA with multiple comparisons. **, *p* < 0.01; ***, *p* < 0.001; ****, *p* < 0.0001. CHX, cycloheximide.

**Figure S10.**
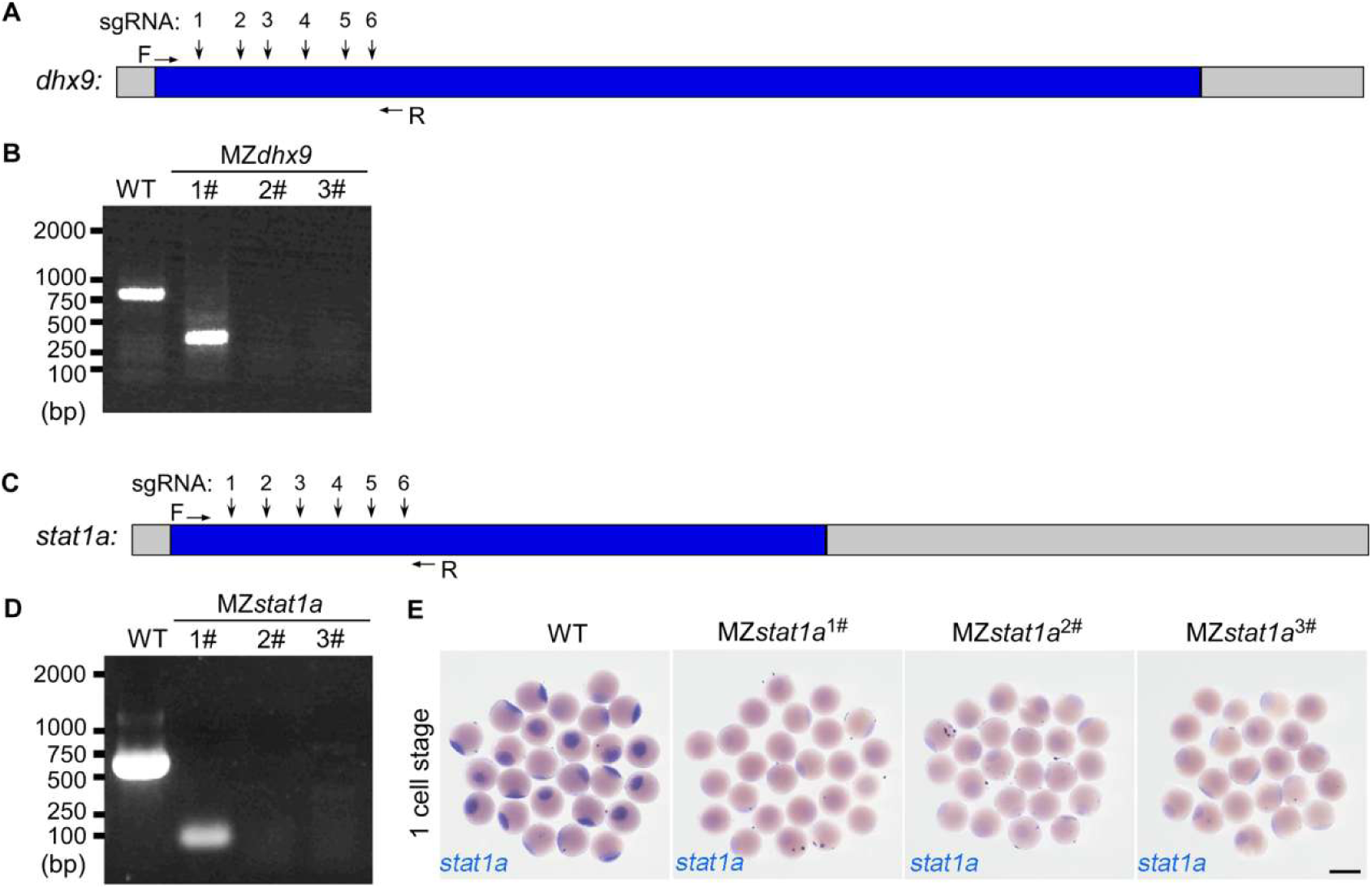
Schematic representation and experimental validation of gRNA-mediated gene knockout of *dhx9* (A, B) and *stat1a* (C-E) in zebrafish. WT, wild type; MZ*dhx9*, maternal and zygotic *dhx9* knockout; MZ*stat1a*, maternal and zygotic *stat1a* knockout. Genotyping for targeting knockout were performed with PCR using genomic DNA as an template (B, D). WMISH assays were performed to examine the expression of *stat1a* in WT and MZ*stat1a* embryos (E). Scale bar, 500 μm.

## Captions for Tables S1 to S4

**Table S1.** The differential expression genes identified in dsRNA transfected mouse ESC and MEF cells, compared to control cells.

**Table S2.** The selected top 100 upregulated genes in dsRNA transfected vs control WT mESCs and their expression in *Prkra* KO, *Dhx9* KO, *p53* KO, *Stat1* KO cells with dsRNA transfection or not.

**Table S3.** The differential expression genes identified in dsRNA transfected human ESC cells, compared to control cells.

**Table S4.** The selected upregulated genes after dsRNA injection in control zebrafish, expression of which were not significant changed in dsRNA injected MZ*tp53* mutant.

